# Unraveling transposable element-mediated regulatory landscape in diverse human immune cells

**DOI:** 10.1101/2024.10.24.620010

**Authors:** Cui Du, Hairui Fan, Jiayao Jiang, Juan Yang, Shuai Chen, Nadezhda E Vorobyeva, Congrui Zhu, Liming Mao, Chenxi Li, Yanhua Li, Wenbin Bao, Ming-an Sun

## Abstract

Transposable elements (TEs) are key contributors to genetic novelty. Despite increasing evidence of their importance, their roles in shaping the regulatory landscape of diverse immune cell populations remain largely unclear. Using single-cell multiome data from human peripheral blood mononuclear cells, we annotated the cell-specific cis-regulatory elements for major immune cell populations and identified a highly cell-specific signature of the overrepresented TE families. Focusing on monocytes that bear fast-evolving transcriptomes, we found that high proportions of their enhancers are TE-derived and bound by multiple pioneer transcription factors. Among them, we confirmed that the core myeloid regulator SPI1 can bind and regulate hundreds of TE-derived enhancers, which further affect the expression of adjacent immune genes. Additionally, interspecies comparison reveals that non-conserved monocyte enhancers are frequently generated by lineage-specific TE insertions, and correlate with the evolved gene expression between human and mouse. Overall, our study supports the importance of TEs in shaping the regulatory landscape of diverse immune cell populations.

## Introduction

Transposable elements (TEs) are mobile DNA elements constituting approximately half of the human genome (*1*). They form two major classes: retrotransposons (Class I) replicate (*i.e*. copy-and-paste) through RNA intermediates to form new insertions, while DNA transposons (Class II) usually move across genome through a “cut-and-paste” mechanism (*2*). Retrotransposons are more abundant than DNA transposons, and they can be further classified as long terminal repeats (LTRs), long interspersed elements (LINEs), and short interspersed elements (SINEs). The sequences of many TEs – particularly the LTRs for Endogenous retroviruses (ERVs) – are rich with transcription factor (TF) binding motifs, thus have an inherent capacity to create cis-regulatory elements (CREs). Further given their fast-evolving nature, TEs are well-regarded as a major source of genetic novelty (*3-5*). To date, there are abundant data linking TEs to mammalian development and diseases (*6-8*). However, how TEs shape the regulatory landscape of diverse immune cells – a highly heterogeneous and fast-evolving system – remains only partially understood.

The mammalian immune system mainly comprises myeloid cells (*e.g*. monocytes, macrophages, and dendritic cells) and lymphocytes (*e.g*. B and T cell subtypes, and natural killer cells), which are differentiated hierarchically from hematopoietic stem cells under the drive of lineage-instructive TFs (*9*). One feature of the immune system is its divergence across species, human populations, or even individuals (*10-13*), likely due to persistent pathogen pressure (*14, 15*). However, most studies on immune evolution focus on gene-level changes (*e.g*. the gain, loss, or mutation of immune genes), with the cis-regulatory mechanisms driving evolved gene expression in diverse immune cell types poorly understood (*16, 17*). However, increasing evidences suggest that the divergence of CREs (particularly enhancers) – rather than protein-coding regions – is the predominant mediator of phenotypic evolution (*18, 19*). With the transcriptomic and epigenomic profiling of diverse immune cells including single-cell sequencing of peripheral blood mononuclear cells (PBMCs) (*20-25*), it provides the opportunity for the investigation of the regulatory mechanisms underlying immune cell evolution.

The regulatory roles of TEs have captured increasing attention in the last decade. Despite numerous studies linking TEs to immunity, most of them focused on the evolution of innate immune response (*26*). For example, a seminar study discovered that primate-specific TEs create thousands of interferon (IFN)-stimulated enhancers to drive innate immunity evolution (*27*). Later studies further uncovered TE-derived enhancers induced by various pathogens or stimuli in human, mouse, and other mammals (*28-31*). The TEs focused by these studies are activated by IFN pathway-related TFs like STAT1/2 and IRF1 (*32*). However, transcriptomic divergence across species is also evident for resting (or unstimulated) immune cells (*17, 33*), and such differences are established during hematopoietic differentiation and may affect immune capacity. Notably, different immune cells show varying degrees of divergence, with macrophages exhibiting more pronounced transcriptomic alterations across species (*34*). Then, could TEs mediate the transcriptomic evolution of resting immune cells? If so, how TEs are associated with the differentiation programs of different immune cells directed by lineage-instructive TFs? Relevant studies are still rare. One previous study found that dozens of TE families (particularly ERVs) are enriched in the enhancers of mouse CD8^+^ T cells, potentially regulating immune gene expression (*35*). In addition, an L1M2a-derived enhancer was reported to regulate the expression of the interferon receptor IFNAR1 in human B cells (*36*). Recently, a study also linked TEs to the tissue-adaptation of different immune cells, particularly regulatory T (Treg) cells (*37*). Despite these advances, the contribution of TEs to the regulatory landscape of diverse human immune cells remains unclear.

Here, we annotated the CREs for diverse immune cell populations by using the single-cell epigenomic data of human PBMCs and identified a highly cell-specific signature regarding the overrepresented TE families. Focusing on monocytes, we found that the core myeloid regulator SPI1 (also known as PU.1) has prevalent binding to TE-derived enhancers and regulates many immune genes. Interspecies comparison further supports the importance of TEs in driving the evolved monocyte gene expression. Our data support the prevalent contribution of TEs in diverse human immune cell populations.

## Results

### Single-cell profiling of CREs in major human immune cell populations

Single-cell sequencing is particularly powerful for analyzing highly heterogeneous immune cell populations (*38*). To identify the CREs in major immune cell populations, we analyzed public single-cell multiome data for human PBMCs (**Data S1**). After pre-processing, we obtained 9,432 cells for single-cell RNA sequencing (scRNA-seq) and 7,161 cells for single-cell assay for transposase-accessible chromatin by sequencing (scATAC-seq). We then annotated 12 major cell populations based on canonical markers (**Fig. 1A, S1**) – with CD14^+^ monocytes the most abundant (**Fig. 1B**). Subsequently, we annotated a total of 105,171 CREs (*i.e*. accessible chromatin regions) for these cell populations, with 33.7% (n=35,443) CREs locate in promoters and most of the remaining CREs occur in intronic or intergenic regions (**Fig. S2**). We further defined cell-specific CREs (csCREs) as those showing significantly higher chromatin accessibility in each major cell population according to the scATAC-seq data. Overall, 28.8% (n=30,297) of the CREs are annotated as csCREs, with their numbers ranging from 1,463 to 11,297 across different immune cell populations (**Fig. 1C, Data S2**).

**Figure 1.**
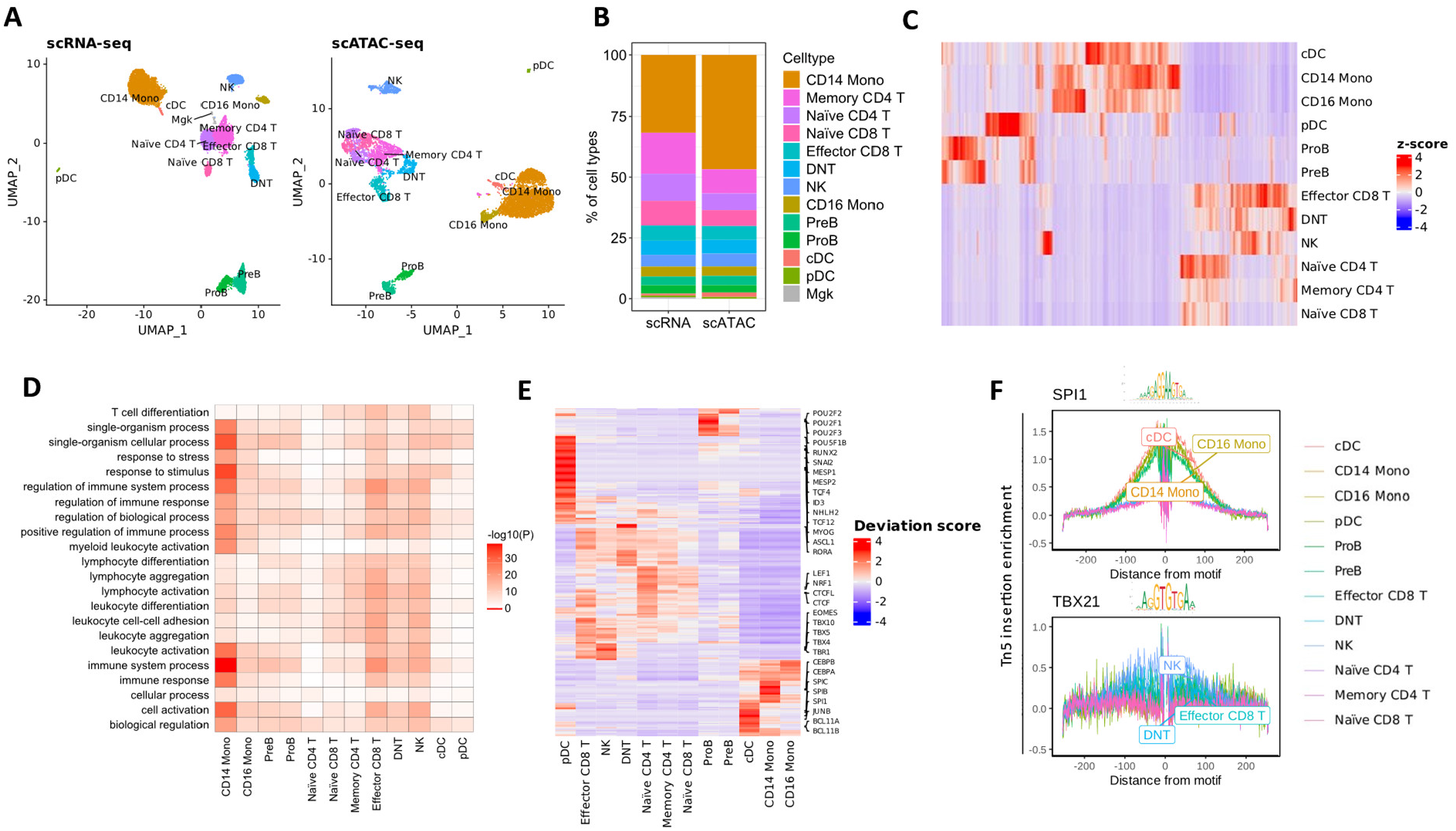
Annotation of the csCREs for diverse immune cell populations in human PBMCs. (**A**) UMAP visualization of different immune cell populations based on the scRNA-seq and scATAC-seq data for human PBMCs. The annotated cell types include CD14^+^ monocyte (CD14 Mono), CD16^+^ monocyte (CD16 Mono), B cell progenitor (ProB), pre-B cell (PreB), natural killer cell (NK), double negative T cell (DNT), conventional dendritic cell (cDC), plasmacytoid dendritic cell (pDC), and megakaryocyte (Mgk). (**B**) Proportions of different immune cell populations based on scRNA-seq and scATAC-seq data, respectively. (**C**) Chromatin accessibility profiles of annotated CREs across various immune cell types. The color gradients represent the column Z-score based on the chromatin accessibility from scATAC-seq data. (**D**) GO enrichment for the csCREs annotated for different immune cell populations. (**E**) Differential TF motifs across different immune cell types. The color gradient represents the bias-corrected deviation scores. The enriched TF motifs for each cell type are labeled on the right side. (**F**) TF footprints for the binding motifs of SPI1 and TBX21 in different types of immune cells.

Next, we determined the functional relevance of the annotated csCREs. As expected, the csCREs for various immune cell types are significantly associated with immune-related functions (**Fig. 1D, Data S3**). For instance, the csCREs for monocytes exhibit high enrichment for immune system processes and immune responses, while those for T cells are enriched for lymphocyte activation and leukocyte differentiation (**Fig. 1D**). Motif analysis was further performed to identify the regulators of these csCREs, with a list of putative upstream TFs uncovered – including many with well-recognized functions in the corresponding immune cell types (**Fig. 1E, S3, Data S4**). For example, we uncover the binding motifs of POU proteins in the csCREs of proB cells (*39*), T-box (*e.g*., EOMES and TBX4/5/21) in NK and T cells (*40, 41*), and SPI1 and CEBP proteins in monocytes (*9*). Notably, the high expression abundance of these TFs usually coincides with the increased accessibility of their binding motifs in the corresponding cell types (**Fig. 1F, S4**). Together, by leveraging the single-cell epigenomic data, we comprehensively annotated and characterized the csCREs for major human immune cell populations.

### Immune csCREs exhibit distinct patterns of overrepresented TE families

Recent studies uncovered the regulatory roles of TEs in mouse T cells and human B cells (*35, 36*), yet a detailed comparison across diverse immune cell populations is still absent. Focusing on the immune csCREs annotated in this study (**Data S3**), we inspected the regulatory involvement of TEs in different human immune cell populations. Given that promoters differ remarkably from other CREs such as enhancers, promoter-proximal and distal csCREs are analyzed separately. We found that most csCREs are distal from promoters (**Fig. 2A**), and as expected, distal CREs show relatively low sequence conservation and high TE overlapping rate (**Fig. S5, 6**). Importantly, we demonstrate that high proportions of distal csCREs are derived from TEs (particularly LTRs/ERVs, LINEs, and SINEs) as annotated by the occurrence of TEs on their center, with their percentages ranging from 20.4% in naïve CD4^+^ T cells to 32.8% in CD16^+^ monocytes (**Fig. 2B**). Notably, the distal CREs for the two subsets of monocytes (CD14^+^ and CD16^+^ monocytes) both show higher overlap rate with TEs relative to other immune cell populations (**Fig. 2B**).

**Figure 2.**
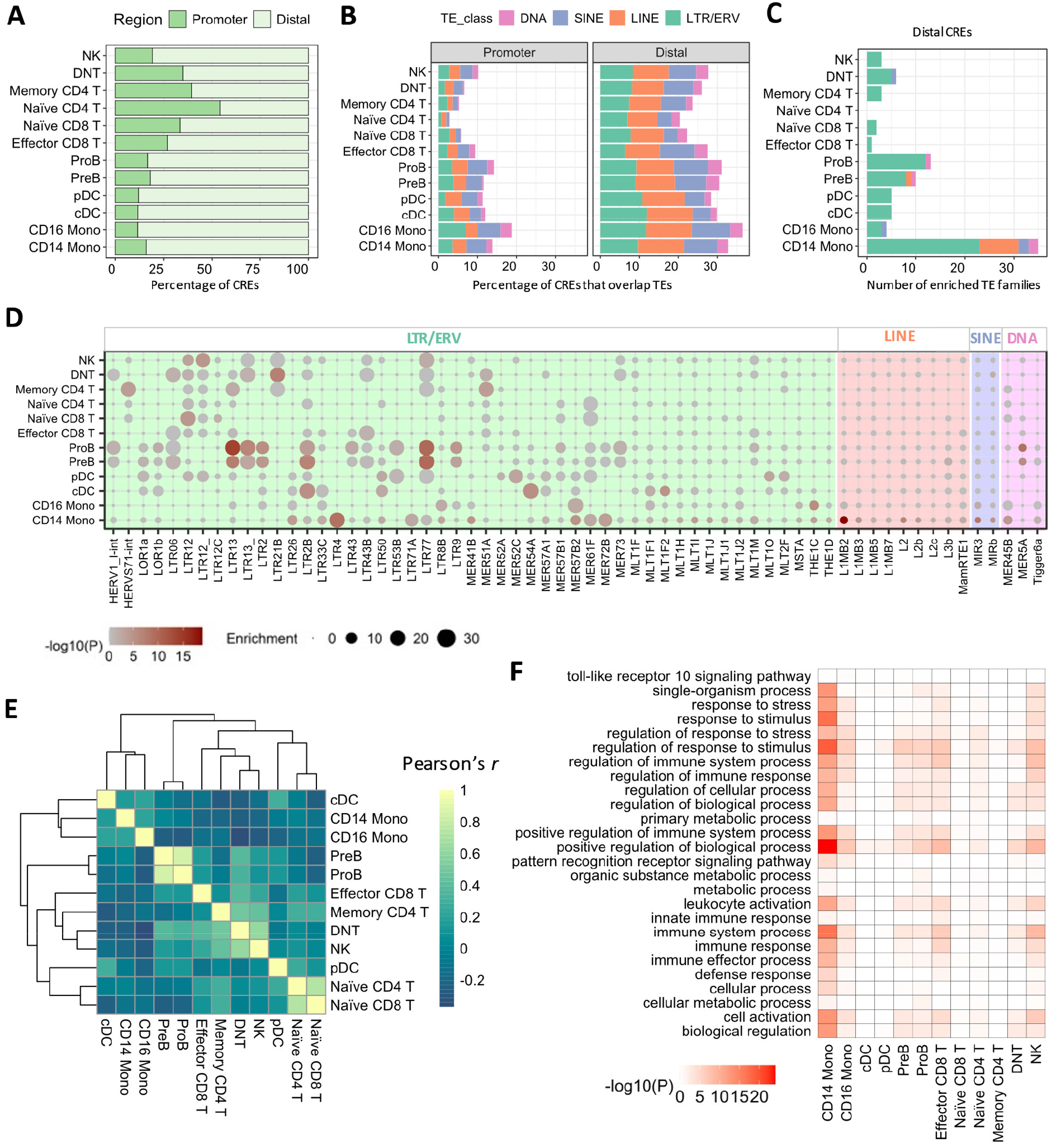
TE enrichment patterns in the csCREs of different human immune cells. (**A**) The proportions of promoter-proximal and distal CREs in different types of immune cells. (**B**) The proportions of csCREs that overlap TEs for different immune cell types. Promoter-proximal and distal csCREs are analyzed separately. **(C)** The numbers of significantly enriched TE families in the distal csCREs for different immune cell types. **(D)** Comparison of the TE enrichment profiles across the csCREs for different immune cell types. All 63 significantly enriched TE families are included for visualization. **(E)** Clustering of different immune cell types based on the TE enrichment profiles of their csCREs. The color gradients represented Pearson’s *r* calculated by using the enrichment fold of different TE families. **(F)** GO enrichment for the distal csCREs of different immune cell types.

We further determined the specific TE families overrepresented in different groups of immune csCREs. In total, we identified 63 TE families significantly enriched in at least one group of distal csCREs, with the majority belonging to LTRs/ERVs (**Fig. 2C, D, Data S5**). In contrast, promoter-proximal csCREs rarely show TE enrichment, except for LTR71B which shows marginal enrichment in cDC cells. Hence, we focused subsequent analyses on distal csCREs. Interestingly, we uncovered evident cell specificity of the TE enrichment signatures. For example, we observed the highest enrichment of LTR4, MER57B2, LTR26, LTR71A and L1MB2 in monocytes, LTR13, LTR2B, LTR77, LTR9 and MER5A in B cells, and LTR12, LTR21B and MER51A in T cells (**Fig. 2D**). Indeed, the cell populations can be clearly clustered based on TE enrichment patterns (**Fig. 2E**). GO analysis demonstrates that TE-derived csCREs are associated with immune-related functions (**Fig. 2F**), further supporting the regulatory involvement of TEs in different immune cell types. Of note, 51.8% (32/63) of the enriched TE families are primate-specific (**Data S6**), thus many of their derived csCREs are newly evolved in primates. Together, TEs form numerous immune csCREs with high cell-specificity, indicating their importance in the regulatory networks of different immune cell populations.

### TE-derived monocyte CREs show enhancer-like epigenetic signature

Monocytes are myeloid-originated cells with crucial roles in innate immunity (*42*). Interestingly, their transcriptomes differ remarkably across species or even among human populations (*17, 34, 43*), implying the fast evolution of their transcriptional regulatory landscape. Further given that TE-derived CREs are poorly studied in monocytes relative to T and B lymphocytes, we focus on this specific cell type for in-depth investigation. To get a better resolution of CRE annotation, we integrated multiple sets of bulk ATAC-seq and epigenomic data of human monocytes (*25, 44, 45*). According to the ChIP-seq data for multiple histone modifications, we demonstrate that distal-CREs often bear high levels of H3K27ac which usually marks active enhancers, and lack the two repressive histone marks H3K27me3 and H3K9me3 (**Fig. 3A**). According to the H3K27ac mark, the majority (91.3%) of distal CREs identified in monocytes are annotated as activate enhancers (**Fig. 3B**).

**Figure 3.**
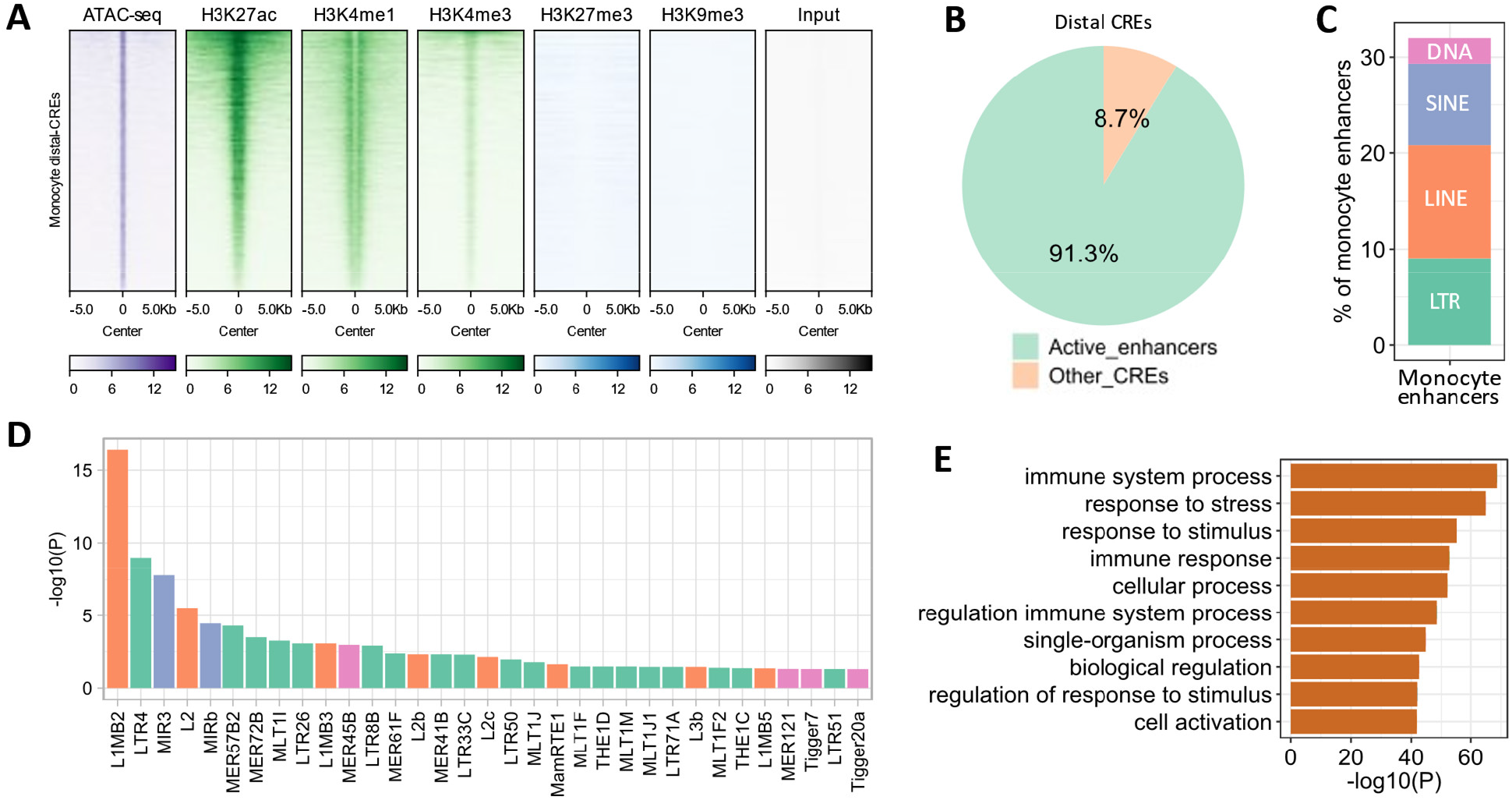
Epigenetic characteristics of human monocyte CREs and their association with TEs. **(A) C**hromatin accessibility and histone modifications (H3K27ac, H3K4me1, H3K4me3, H3K27me3, and H3K9me3) for the distal CREs in human monocytes. This figure is based on the ATAC-seq and ChIP-seq data for purified human monocytes. **(B)** The proportion of monocyte distal CREs that are annotated as active enhancers based on H3K27ac data or not. **(C)** The proportions of monocyte enhancers that overlap major TE classes, including LTR, LINE, SINE and DNA transposons. **(D)** The TE families that are significantly enriched in monocyte enhancers. Different classes of TEs are indicated by color. **(E)** GO enrichment for the monocyte TE-enhancers. The top ten enriched GO terms are shown.

Focusing on the epigenetically-annotated monocyte enhancers based on H3K27ac mark, we found that about one-third of them are derived from TEs (**Fig. 3C**), which are thereby denoted as TE-derived enhancers (abbreviated as TE-enhancers). In total, we identified 33 TE families significantly enriched in monocyte enhancers (**Fig. 3D, S7, Data S7**), with 84.8% (n=28) matching those identified in the csCREs for CD14^+^ monocytes (**Data S5**). While most of these enriched TE families belong to LTR/ERV (*e.g*. LTR4 and MER57B2), we also noticed several families of LINE (*e.g*. L1MB2), SINE (*e.g*. MIR3) and DNA transposons (*e.g*. MER45B). As expected, monocyte TE-enhancers are also highly associated with myeloid and immune-related functions (**Fig. 3E**). Further inspection revealed the presence of TE-enhancers near multiple important immune genes, such as *LYN, TRAF3, cGAS* and *MYD88* (**Data S8**). Of note, *cGAS* and *MYD88* are key genes of the cGAS/STING pathway, which mediates the sensing of intracellular double-stranded DNA molecules (*46*). These data suggest that TEs form numerous active enhancers in monocytes, which may influence the expression of adjacent immune genes.

### Screening the putative regulators of TE-derived monocyte enhancers

After uncovering the enrichment of TEs in monocyte enhancers, we further screened for the TFs that may bind and regulate TE-derived monocyte enhancers. By motif analysis in monocyte TE-enhancers, we uncovered 30 candidate TFs (**Data S9**), which largely match the differential TF motifs identified by using chromVAR across different immune cell types (**Fig. 1E**). Further, several of these TFs are known to play critical roles in monocytes. For example, SPI1 is a key regulator for monocyte development and differentiation, and ATF4, CEBPA/B, IRF1, and STAT1/3 are also important for monocytes (*47-49*). Importantly, several of these TFs (particularly SPI1 and CEBPA/B) have relatively high expression in monocytes compared to other immune cell types (**Fig. S4**). Therefore, these TFs are considered putative upstream regulators of TE-enhancers in monocytes.

We then examined the binding of these TFs in human monocytes based on public ChIP-seq data, focusing on eight TFs including SPI1, CEBPA/B, ATF4, IRF1, SREBF2, and STAT1/3 (**Data S1**). Most of the binding peaks for these TFs reside outside of promoter regions and show strong H3K27ac mark, indicating they predominantly correspond to active enhancers (**Fig. 4A, C, S8**). We found that large proportions of their distal peaks overlap TEs (primarily LTRs, LINEs, and SINEs), ranging from 22.3% for STAT3 to 42.0% for CEBPA (**Fig. 4B**). Among the TEs bound by these TFs, LINE elements have the highest proportion. Specifically, the peaks of these TFs are significantly enriched with dozens of TE families, with the majority belonging to LTRs and LINEs (**Data S10**). Of note, many of the 33 TE families significantly enriched in monocyte enhancers are also frequently bound by the analyzed TFs (*e.g*. L1MB2/3/5 for CEBPs, and LTR50, MER57B2, L2/L2b/L2c for SPI1) (**Fig. 4D**). In particular, 90.9% of the TE families enriched in monocyte enhancers also significantly overlap SPI1 peaks, which is the highest in these TFs (**Fig. 4E**) – implying the pivotal role of SPI1 in regulation monocyte TE-enhancers. We further inspected the MER57 subfamilies (MER57A1/B1/B2) which are significantly enriched in SPI1 peaks, and confirmed that most of them harbor SPI1 binding motif (**Fig. 4F**). Overall, we identified a set of TFs which are putative regulators of TE-enhancers in monocytes.

**Figure 4.**
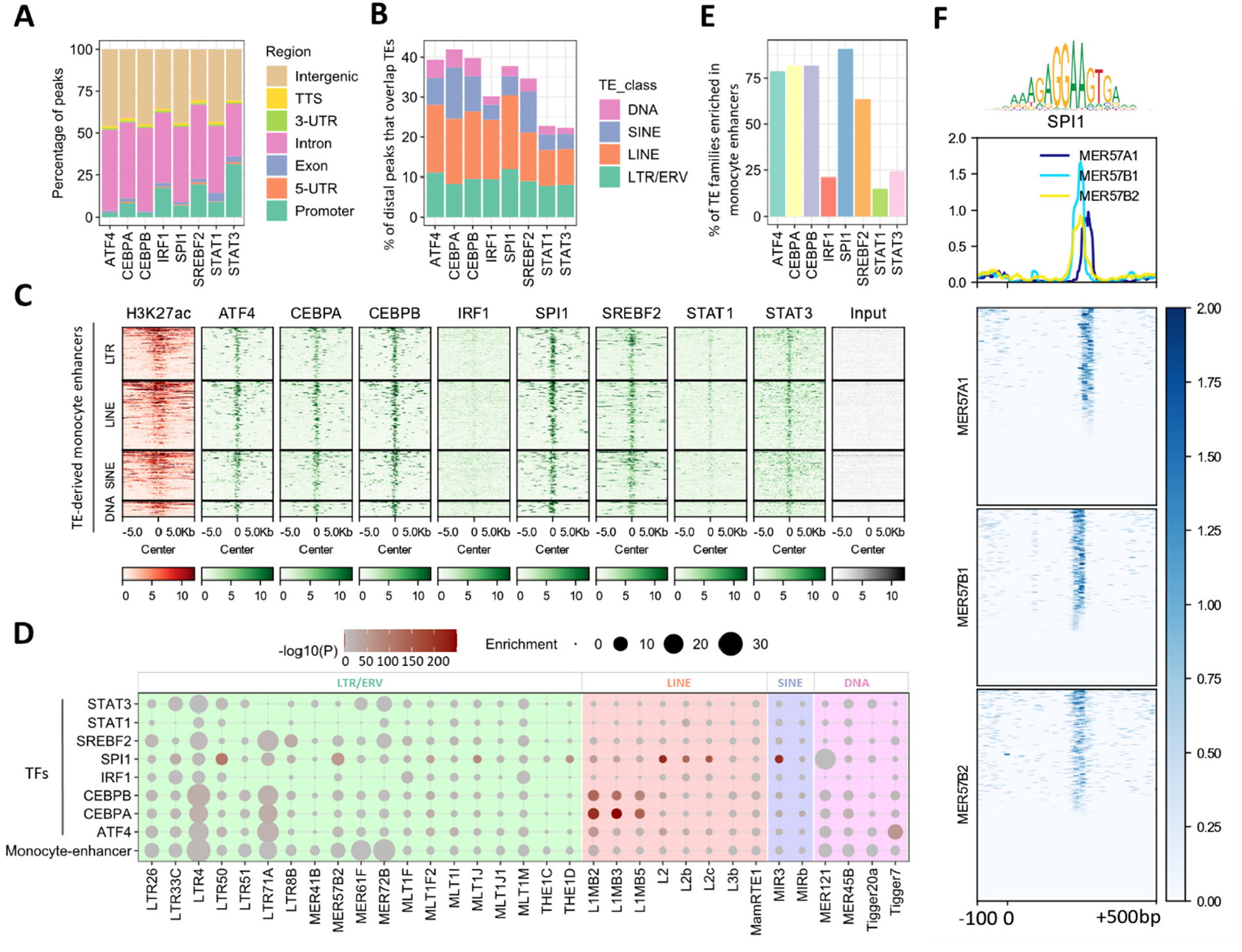
Inferrence of the TFs that may regulate TE-derived enhancers in human monocytes. **(A)** Genomic distribution of the 8 TFs which may regulate TE-derived enhancers in human monocytes. **(B)** The proportions of the distal peaks of the 8 TFs that overlap TE-derived monocyte enhancers. **(C)** The epigenetic characteristics of TE-enhancer binding to four TE classes in putative regulatory factors of TE-derived monocyte enhancers. **(D)** TE enrichment profiles in monocyte enhancers and the binding peaks of different TFs. The 33 TE families enriched in monocyte enhancers are included for visualization. **(E)** Comparison of the TE families enriched in monocyte enhancers relative to the binding peaks of each TF. The y-axis shows the percentage of TE families enriched in monocyte enhancers that are also enriched in the peaks of each TF. **(F)** Schematic representation of the SPI1 motif within the sequences of different MER57 subfamilies (MER57A1/B1/B2), which are significantly enriched in SPI1 peaks. The color gradients represent the motif matches based on the log-likelihood ratio.

### Regulation of TE-derived enhancers and immune genes by SPI1 in monocytes

SPI1 is a pioneer TF that has been extensively studied in monocyte and other myeloid lineages (*47, 48, 50*). After revealing its prevalent binding on TE-derived monocyte enhancers, we further determined its regulatory function by performing siRNA-mediated knockdown in THP-1 monocyte cells. The knockdown efficiency was confirmed by both qRT-PCR and Western blotting (**Fig. 5A, B**). We first examined the transcriptomic alterations after SPI1 knockdown (SPI1KD) using RNA-seq (**Fig. S9**). In total, we identified 1,076 differentially expressed genes (DEGs) after SPI1KD, with 510 up-regulated and 566 down-regulated (**Fig. 5C, Data S11**). Interestingly, the down-regulated genes are tightly associated with immune-related functions (**Fig. 5D**), suggesting the critical role of SPI1 for immune gene expression. We then performed H3K27ac ChIP-seq to compare the activation status of SPI1-bound loci before and after SPI1KD. A total of 4,154 genomic loci show altered H3K27ac level (**Fig. 5E, Data S12**), and 36.4% (n=1,510) of them are bound by SPI1 (**Fig. 5G**). These loci are putative target loci under the direct regulation of SPI1. We further compared the loci with increased or decreased H3K27ac levels after SPI1KD and found that significantly higher percentage of H3K27ac-decreased loci are originate bound by SPI1 (**Fig. 5F, S10**). These data suggest that SPI1 prefers to activate its target loci, which agrees with a previous study on the predominant role of SPI1 for transcriptional activation (*51*).

**Figure 5.**
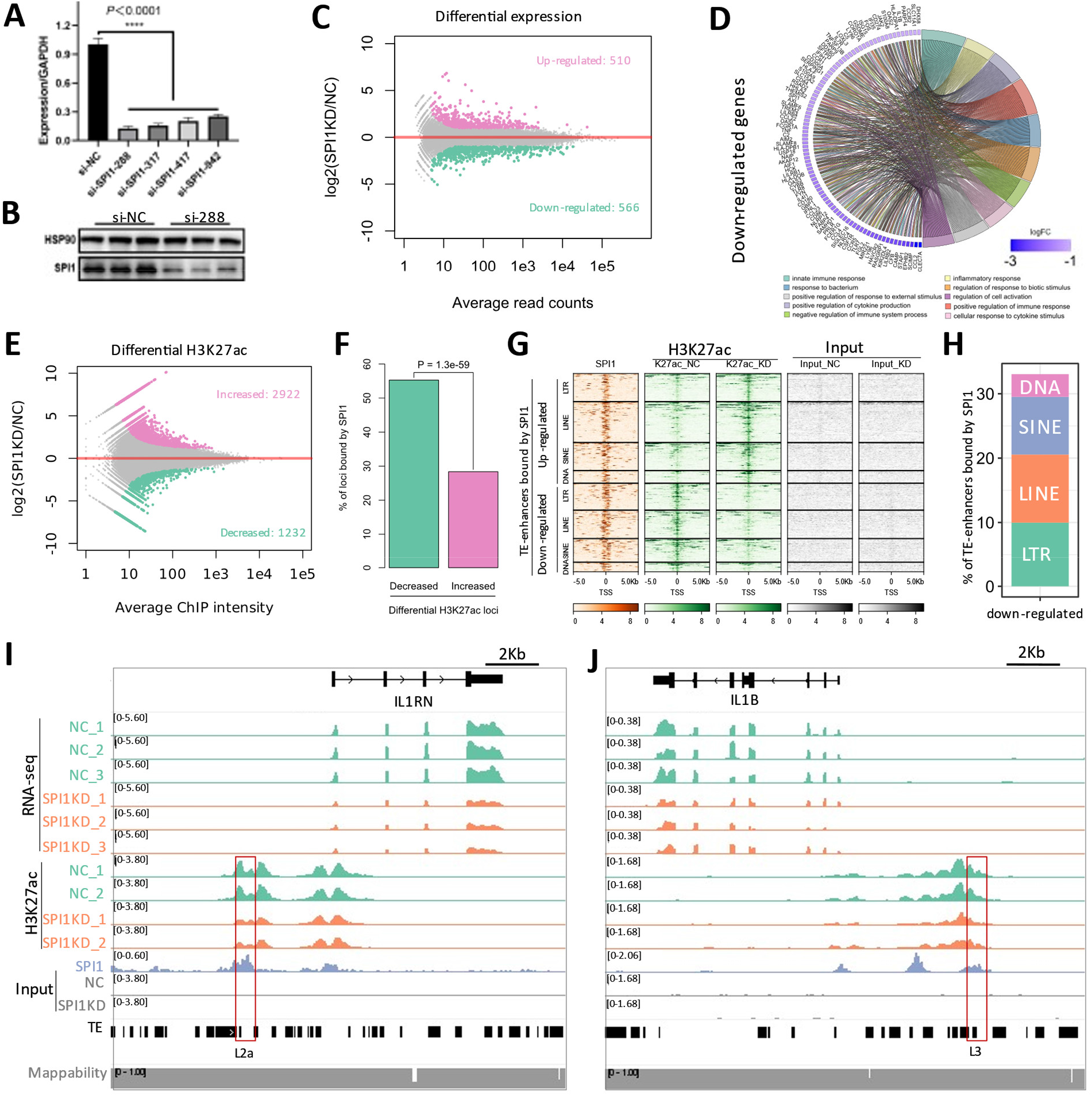
Regulation of TE-enhancers and immune genes in human monocytes by SPI1. **(A, B)** Validation of the siRNA interference efficiency of SPI1 in THP-1 monocyte cells by using qRT-PCR and Western blot. **(C)** Differential gene expression after SPI1KD based on RNA-seq data. **(D)** GO enrichment of the down-regulated genes after SPI1KD. **(E)** Differential H3K27ac intensity after SPI1KD based on ChIP-seq data. **(F)** Comparison of up-regulated vs. down-regulated H3K27ac loci regarding their overlap to SPI1 peaks. P-value based on Fisher’s Exact Test is indicated. **(G)** Epigenetic characteristics of SPI1-bound TE-enhancers that show reduced H3K27ac levels after SPI1KD. **(H)** The proportions of down-regulated enhancers bound by SPI1 overlap four major classes of TEs. **(I, J)** IGV tracks show the SPI1-bound TE-derived enhancers upstream of IL1RN and IL1B, respectively. The reduced activity (as reflected by decreased H3K27ac level) of the TE-derived enhancers correlated with the decreased expression of adjacent genes.

We further investigated whether SPI1 regulates the expression of some immune genes in monocytes through TE-derived enhancers. Focusing on the 681 SPI1-bound loci showing decreased H3K27ac level after SPI1KD, we found that 33.9% (n=231) of them are derived from TEs (**Fig. 5H**). Moreover, dozens of TE-derived loci are adjacent to genes down-regulated after SPI1KD (**Data S11**). For example, an L2b element located about 2 kb upstream of the Interleukin 1 receptor antagonist (*IL1RN*), which is a crucial immune gene expressed in monocytes (*52*). The H3K27ac level of the SPI1-bound L2b element and the expression level of *IL1RN* are concurrently decreased after SPI1KD (**Fig. 5I)**, indicating that this L2b element may mediate the expression of *IL1RN* under the control of SPI1. SPI1 also binds an L3-derived enhancer to control the expression of interleukin 1B (*IL1B*) (**Fig. 5J**), which is a crucial monocyte-secreted cytokine known to be under SPI1-mediated transcriptional regulation (*53-55*). Apart from these two genes, we also identified dozens of immune genes (*e.g. CCR2, CYBB*, and *MSR1*) likely under the control of SPI1-regulated TE-enhancers (**Data S13**). Taken together, these results indicate that SPI1 regulates the expression of many monocyte genes through its binding to TE-derived enhancers.

### Divergence of TE-derived monocyte enhancers between human and mouse

Since many TE families overrepresented in human monocyte enhancers are primate-specific (**Data S6**), we wonder if TE-enhancers can drive the divergence of monocyte gene expression across species. To answer this, we compared the epigenomic and transcriptomic data between human and mouse monocytes (**Data S1**). We first used H3K27ac ChIP-seq data to annotate monocyte enhancers, and found that high proportions of monocyte enhancers are not conserved across species (**Fig. 6A**). Impressively, TEs frequently overlap lineage-specific monocyte enhancers in both species, notably 51.7% for human and 43.3% for mouse (**Fig. 6B**). Of note, LINEs show remarkably higher overlap rate to monocyte enhancers in human relative to mouse (**Fig. 6B**), despite that the global abundance of LINEs is comparable (21.7% *vs*. 20.0%) in both genomes. In contrast to human-mouse shared enhancers which only have 7 significantly enriched TE families (**Fig. S11**), as many as 69 TE families are significantly enriched in lineage-specific enhancers (**Fig. 6C, Data S14**). Importantly, many of the enriched TE families are species-specific, such as the primate-specific LTRs (LTR4, LTR26, LTR8B, LTR36, MER72, HERV3-int), and rodent-specific SINEs (ID_B1, B3, B3A, B4A) and LTRs/ERVs (MTD, MTEa, ORR1D2, ORR1E). These data suggest the importance of TEs for monocyte enhancer evolution in human and mouse.

**Figure 6.**
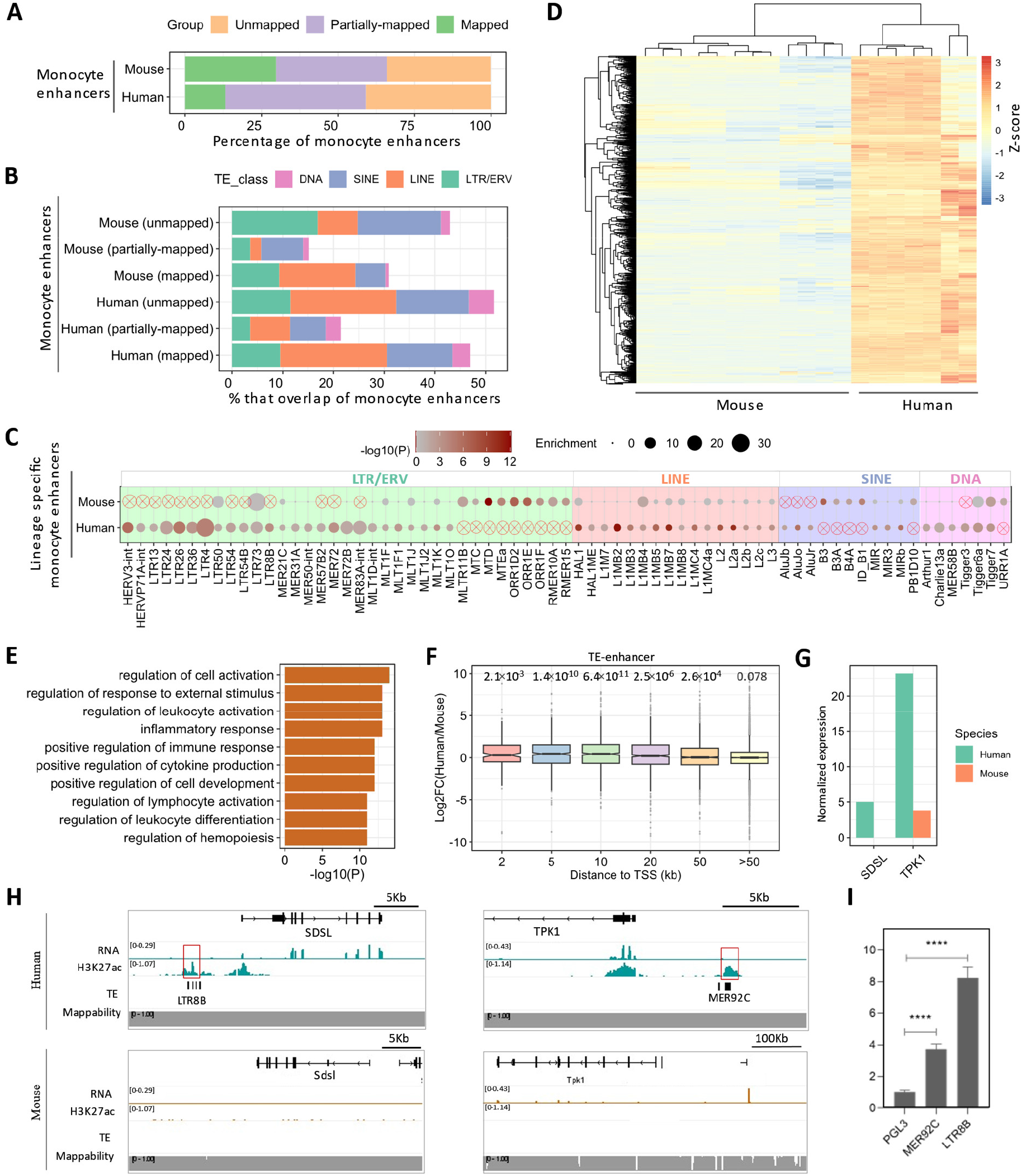
Contribution of TEs to the transcriptomic divergence between human and mouse monocytes. **(A)** The proportion of monocyte enhancers in human and mouse that are annotated as conserved or species-specific. **(B)** The overlap of TEs with species-specific or conserved monocyte enhancers in human and mouse. **(C)** TE enrichment profiles in lineage-specific monocyte enhancers for human and mouse. The primate- or rodent-specific TE families are marked with ⊗. **(D)** Expression profiles of the genes with higher expression in human relative to mouse. The color gradient represents the Z-score calculated from normalized TPM values. **(E)** GO enrichment results of the genes with higher expression in human. **(F**) Interspecies expression alterations of the genes associated with lineage-specific monocyte enhancers derived from TEs. The boxplots compare the genes with human-specific monocyte enhancers occurring at different distance thresholds. P-values calculated by using Student’s t-test again zero are denoted. **(G**) Expression level of *SDSL* and *TPK1* between human and mouse monocytes, as measured by normalized TPM values. **(H**) IGV tracks show the transcriptomic and epigenomic pattern flanking *SDSL* and *TPK1*. **(I)** Luciferase reporter assay results for the TE elements associated with the putative lineage-specific enhancers adjacent to *SDSL* and *TPK1*, respectively.

We further determined the transcriptomic alteration between human and mouse monocytes following a previous study (*56*). A total of 2,188 interspecies DEGs are identified, with 1,083 and 1,105 showing higher expression in human and mouse, respectively (**Fig. 6D, S12A, Data S15**). The DEGs with higher expression in either human or mouse monocytes are both associated with defense response (**Fig. 6E, S12B**). Indeed, many DEGs are canonical immune genes, such as *LYZ, S100A8/9, CD14, ITGB2, TNFSF13B* in human and *Ly6e, Slpi, Zbp1* in mouse (**Data S15**). To determine whether TEs mediate the interspecies expression divergence, we inspected the genes adjacent to lineage-specific enhancers, and found they indeed exhibit increased expression in human (**Fig. 6F, S13**). We identified hundreds of TE-enhancer adjacent genes showing higher expression in human monocytes (**Data S16**), and then performed a luciferase reporter assay in K562 myeloid cell line to assess the enhancer activity of three TE elements, which tend to form lineage-specific enhancers adjacent to *SDSL, TPK1* and *NOPCHAP1*, respectively. All these genes exhibit significantly greater expression in human monocytes relative to mouse (**Fig. 6G-I, S14**), and SDSL shows myeloid-enriched expression (**Fig. S15**) according to Human Protein Atlas (HPA) database (*57*). Our data confirmed the strong enhancer activity for two out of the three candidate TE elements (**Fig. 6J**). One is the LTR8B element upstream of *SDSL*, which is a metabolism-related gene associated with mitochondrial dysfunction and acute myeloid leukemia (*58, 59*). The other is the MER92C element upstream of *TPK1*, which is a protein kinase mediating the cell response to environmental stress (*60, 61*). Surprisingly, the MLT1F2 element upstream of *NOPCHAP1* shows a repressive function (**Fig. S14B**), indicating the dependence on other DNA elements. Together, our data suggest that TEs create numerous species-specific monocyte enhancers in both human and mouse, which may contribute to the interspecies transcriptomic divergence of monocytes.

## Discussion

A remarkable signature of the immune system is its fast evolution, as evidenced by its divergence across species, human populations, and even individuals. It has long been recognized that pathogen pressure is the major extrinsic driver of immune evolution – as conceptually proposed by the “Red Queen” hypothesis and then further refined by later studies (*12, 14, 62*). Meanwhile, the intrinsic genetic determinants underlying immune evolution also attracted increasing studies. TEs are regarded as a major source of genetic innovation (*1, 8*). In the past decade, the importance of TEs during mammalian development has been extensively studied, with particular attention on embryonic stem cells (ESCs) and placentae (*6, 56, 63, 64*). The importance of TEs for immunity has also long been recognized since the emergence of adaptive immunity in jawed vertebrates depends on the two TE-derived RAG1/2 transposases (*65*). Nevertheless, the cis-regulatory roles of TEs in diverse immune cells are still poorly understood – partly due to the extreme cellular complexity of the immune system. Here, we characterized the regulatory roles of TEs in diverse human immune cells, first achieved a global picture by using single-cell sequencing data, and then focused on monocytes to elucidate their functions and underlying mechanisms. Unlike most previous studies on innate immune response (*28-31*), this study focused on resting immune cells which highlights the roles of TEs for immune cell differentiation.

Single-cell sequencing is particularly powerful for studying the highly heterogeneous immune system (*38, 66*). By integrating the scATAC-seq data of human PBMCs, we annotated the CREs for major immune cell populations, including different myeloid and lymphocytic subsets. It is worth noting that, a recent study used scATAC-seq to analyze tissue-residing immune cells (particularly Tregs) and linked TEs to their tissue adaptation (*37*). Focused on the annotated immune csCREs, we confirmed their association with the corresponding lineage-instructive TFs, such as SPI1, IRF1, and CEBP in monocytes, GATA3, TBX21, TCF7 in T cells, and SPI1, PAX5 in B cells (*9, 67*). Interestingly, dozens of TE families are enriched in these CREs with high cell-specificity. For example, we noticed many TE families exclusively enriched in the CREs for monocytes (*e.g*. LTR4, MER57B2, LTR71A), B cells (*e.g*. LTR9, LTR2, LTR43) or T cells (*e.g*. LTR12, MER51A). Further, the TE families enriched within immune CREs differ remarkably from those for human ESCs and placentae (*56, 63, 64*). These data suggest that many TE families are selectively adopted by distinct immune cell lineages. Accompanying recent studies linking TEs to B and/or T lymphocytes (*35-37*), our study provides a global profile of the TE regulatory landscape across various immune cell types and supports the widespread participation of TEs in the regulatory landscape of different immune cells.

Among the profiled immune cells, we performed an in-depth analysis of monocytes, which are crucial for host immunity and tissue homeostasis (*42*). Despite previous research on the TE-mediated IFN response in monocytes (*27*), how TEs shape the regulatory landscape of resting monocytes remains largely unclear. By analyzing CREs identified by scATAC-seq, we identified many TE families exclusively enriched in monocytes relative to other immune cell types – suggesting that these TEs are selectively activated in monocytes. In addition to LTRs/ERVs, we surprisingly found that other classes of TEs – particularly those belonging to LINEs (*e.g*. L1MB2, L1MB3, L2, L2b) – are also highly enriched in monocyte CREs with high cell specificity. This observation is further confirmed by the analysis of monocyte enhancers as annotated by the H3K27ac mark. LINEs constitute over 20% of the human genome, with LINE-1 (L1) known as the only type of TEs with autonomous transposition capacity (*1, 6*). Unlike LTRs with inherited functions as CREs, most previous studies on LINEs focus on their *trans* function by their produced RNA or proteins, with their cis-regulatory roles largely unexplored. A recent study provides the first experimental evidence that specific LINE1 elements (L1Md_Ts) can form enhancers in mouse ESCs (*68*), and other studies suggest that LINEs can also form promoters or insulators (*69, 70*). Our study demonstrates that LINEs also form numerous enhancers in human monocytes. Surprisingly, even though most of the LINE families (*e.g*. L2b and L1MB2) enriched in human monocyte enhancers are also owned by mice, LINEs tend to create remarkably higher amounts of monocyte enhancers in human relative to mouse. These findings suggest that lineage-specific insertions of ancient-originated LINE elements – together with LTRs/ERVs including some primate-specific ones – are crucial in shaping the regulatory landscape of human monocytes.

Unlike in ESCs and placentae, where the TE activators have been extensively studied (*63, 64, 71, 72*), the TFs underlying TE activation in monocytes are unclear. Through computational screening, we identified multiple TFs that frequently bind TE-enhancers in monocytes, including several that have been known to be important for monocyte differentiation or related myeloid lineages (*73, 74*). SPI1 was selected for in-depth analysis because it is a master pioneer TF that mediates the late specification of monocytes and derived macrophages (*48, 50, 75*). Indeed, the ectopic expression of SPI1 alone can induce a myeloid gene program (*51, 76*). We found that many immune genes are down-regulated after SPI1KD, suggesting the critical function of SPI1 in monocytes. Our data indicate that SPI1 mainly acts as a transcriptional activator, since a higher proportion of genomic loci with decreased H3K27ac after SPI1KD are originally bound by SPI1. Meanwhile, there are also many SPI-bound loci with unchanged or even decreased H3K27ac after SPI1KD, implying that the function of SPI1 across the genome can be context-dependent. Several other TFs (*e.g*. CEBPs and IRF1) frequently co-bind with SPI1 across the genome. This is probably because these TFs have combinatorial functions over some targeted loci, as recent evidence reveals that SPI1 and CEBPA collaborate to open closed chromatin regions (*77*). Regarding the TE binding profiles, the screened TFs show both similarities and differences regarding the enriched families – which is not surprising given their co-binding on many genomic loci. Relative to other TFs, SPI1 shows a higher degree of enriched on several families of LTRs (LTR50, MER57B2, MLT1J, THE1D), LINEs (L2, L2b, L2c) and SINEs (MIR3). Inspection of representative TE families (MER57A1/B1/B2) uncovered the presence of SPI1 motifs in their sequence, thus explaining why SPI1 can recognize and bind these TEs. By integrative comparison of the transcriptomic and epigenomic data, we found that SPI1 tends to regulate certain genes through its binding on TE-derived enhancers. Based on these results, we propose that under the regulation of core TFs like SPI1, TEs participate in the monocyte regulatory network to regulate the expression of specific immune genes.

To understand the evolutionary divergence of monocytes, we performed transcriptomic and epigenomic comparisons between human and mouse. As expected, hundreds of genes show differential expression across species, and many of them are immune-related. Correspondingly, we found that high proportions (>30%) of monocyte enhancers are not shared across species, and about 50% of species-specific monocyte enhancers are TE-derived – suggesting the importance of TEs in driving monocyte regulatory evolution. Correlation analysis indicates that TE-derived enhancers may influence the evolved expression of some adjacent genes, and luciferase reporter assay further confirmed the enhancer activity of representative TE elements associated with two genes showing higher expression in the monocytes of human relative to mouse. We identified dozens of TE families significantly overrepresented in non-conserved monocyte enhancers, which show several intriguing patterns: 1) Most enriched LTR families are primate- or rodent-specific, suggesting the role of young LTR families in creating novel monocyte enhancers; 2) Multiple ancient LINE families show a higher degree of enrichment in human, suggesting they create more primate-specific insertions to form monocyte enhancers; 3) Higher proportions of monocyte enhancers overlap SINEs in mouse relative to human, and more SINE families (most are rodent-specific) are significantly enriched in mouse-specific monocyte enhancers. Overall, these data suggest that TEs have species-biased functions in shaping the regulatory landscape of human and mouse monocytes, and novel monocyte enhancers are not only created by species-specific LTR/SINE families but also lineage-specific insertions of TEs belonging to ancient LINE families.

Overall, this study represents a comprehensive investigation of the TEs landscape across various immune cell types and highlights the cis-regulatory function of TE-derived enhancers in monocytes and other immune cell populations.

## Materials and Methods

### Cell culture

THP-1 monocyte cell line was purchased from Procell (Wuhan, China). The culture medium consists of RPMI-1640 supplemented with 10% fetal bovine serum (FBS), 0.05 mM β-mercaptoethanol, and 1% penicillin/streptomycin (P/S). K562 cells (human myeloid leukemia cell) are kindly provided by Dr. Shaobin Shang, and the culture medium consists of RPMI-1640 supplemented with 10% (FBS) and 1% penicillin/streptomycin (P/S). Cells were grown in a constant-temperature incubator at 37°C with 5% CO_2_.

### SPI1 knockdown

We performed siRNA-mediated knockdown to disrupt SPI1 expression. The siRNA sequences are summarized in **Data S17**. The efficiency of the siRNA experiment was validated by qRT-PCR and Western blot. The detailed experimental procedures for related experiments are provided in **SI Appendix, Materials and Methods**.

### Luciferase reporter assay

The sequences of selected TE elements (**Data S17**) were synthesized and inserted into pGL3-basic vectors (GeneCreate, Wuhan, China). K562 cells were co-transfected with pGL3-basic (containing TE sequence or blank control) and pRL-TK vectors. Luciferase activities were measured with the Dual-Luciferase Reporter Assay System (Vazyme, DL101, Nanjing, China) 24h after transfection. Data for firefly luciferase activity were normalized to Renilla luciferase activity. All experiments were performed at least three times independently.

### ChIP-seq

ChIP-seq was performed following a previous study (*56*), with the detailed experimental procedure provided in **SI Appendix, Materials and Methods**. Two biological replicates were used per sample. The H3K27ac antibody is from SantaCruz (sc-390405). ChIP-seq libraries were constructed using Takara SMARTer ThruPLEX DNA-seq Kit (QIAGEN, Cat# 28004) and sequenced as 100 bp paired-end reads with the DNBSEQ (MGI) platform.

Raw reads were trimmed with Trim Galore! v0.6.4 (*78*) and then aligned to the corresponding reference genome (GRCh38 for human, GRCm38 for mouse) using Bowtie2 v2.3.5 (*79*) with default settings. PCR duplicates were removed using the *rmdup* function of SAMtools v1.13 (*80*). After confirming the data reproducibility, reads from biological replicates were pooled together for further analysis. Peak calling was performed with MACS v2.2.6 (*81*) with settings: --g hs --q 0.05. The peaks were further cleaned by removing those that overlap ENCODE Blacklist V2 regions (*82*). Differential binding analysis was performed using DiffBind v3.4.11 (*83*) with settings: minOverlap = 1, summits = 250, method = DBA_EDGER.

### ATAC-seq

ATAC-seq data are collected from public studies (**Data S1**). The ATAC-seq data were analyzed following the same procedure as the ChIP-seq data.

### RNA-seq

Detailed experimental procedure for RNA-seq is provided in **SI Appendix, Materials and Methods**. Total RNA was isolated from three biological replicates using TRIzol (Qiagen), and RNA integrity and concentration were assessed by the Qubit™ RNA HS Assay (Invitrogen, Q32852). RNA-Seq libraries were sequenced as 150 bp paired-end reads with the DNBSEQ (MGI) platform.

Raw RNA-seq reads were trimmed with Trim Galore v0.6.4 (*78*). Transcript per million (TPM) values were calculated with RSEM v1.3.2 (*84*). We aligned trimmed reads to the reference genome of human (GRCh38) or mouse (GRCm38) using STAR v2.7.3(*85*), and then obtained gene-level read counts by using the *featureCount* function from subread v2.0.0 (*86*). At last, DEGs were identified by using DESeq2 v1.30.1 (*87*) with the cutoff: FDR < 0.05, |log2foldChange| > 1.

### scRNA-seq

Raw scRNA-seq data for human PBMCs were downloaded from 10 × Genomics website (**Data S1**). Cell Ranger (10 × Genomics) was used for demultiplexing, followed by read alignment to human reference genome (GRCh38). Seurat (*88*) was used for data filtering, normalization, dimensionality reduction, and clustering processes. Low-quality cells were filtered out based on the criteria: gene count > 200, gene count < 20,000, mitochondrial content < 20%.

### scATAC-seq

Raw scATAC-seq data for human PBMCs were downloaded from 10 × Genomics website (**Data S1**). Cell Ranger ATAC v2.1.0 (10 × Genomics) was used for demultiplexing, followed by read alignment to human reference genome (GRCh38). Signac v2.1.0 (*89*) was used for subsequent analysis. Low-quality cells were removed based on the criteria: peak region fragments > 3000, peak region fragments < 30,000, pct_reads_in_peaks > 15%, blacklist ratio < 0.05, nucleosome signal < 4, TSS enrichment > 3. After filtering, a total of 110,202 peaks from 8,379 cells remained for latent semantic indexing (LSI) analysis (*90*). Clustering and dimensionality reduction were then performed on the corrected LSI components. Finally, we obtained 16 clusters with resolution = 0.5 and dims = 2:30. The activity of each gene was quantified by examining the local chromatin accessibility, including the gene body and the 2 kb region upstream of TSS. The accessible chromatin peaks for each cell type were identified by using MACS2 (*81*). Differential chromatin accessibility analysis was performed by using the *FindAllMarkers* function. Differentially accessible chromatin regions were identified with the threshold: Bonferroni-adjusted P-value < 0.05, log2foldchange > 0.25. The genomic regions containing accessible chromatin peaks were annotated by ChIPSeeker v1.36.0 (*91*).

### Integration of scRNA-seq and scATAC-seq data

To better interpret the scATAC-seq data, we classify cells based on a scRNA-seq experiment for matched human PBMC samples. We utilize methods for cross-modality integration and label transfer. The shared correlation patterns between scATAC-seq gene activity and scRNA-seq gene expression were identified by the *FindTransferAnchors* function (reduction = cca). Then the cell type label of each cell in scATAC-seq data was predicted by the *TransferData* function (weight reduction = lsi, dim = 2:30). A total of 7,161 cells were retained after filtering using a maximum prediction score ≥ 0.5.

### Transcription factor motif analysis

Single-cell TF motif activity was estimated for a set of 849 TFs from the Catalog of Inferred Sequence Binding Preferences (CIS-BP) database (human_pwms_v2) using the *RunChromVAR* function in Signac v2.1.0 (*89, 92*). Differential TF activity between cell types was calculated by the *FindMarkers* function with threshold: |log2foldChange| > 1, Bonferroni-adjusted P < 0.05. The occurrence of the SPI1 motif was identified by using FIMO v5.4.1 (*93*) with settings: --text --thresh 1E-4. The position-weight matrices for SPI1 were obtained from the JASPAR database (*94*). DeepTools (*95*) was used to plot over MER57A1/B1/B2 elements as heatmaps.

### Reference genome and annotation

Reference genome and gene annotation for human (GRCh38) and mouse (GRCm38) were downloaded from the Ensembl database (release 102) (*96*). TE annotations were downloaded from the RepeatMasker website (http://www.repeatmasker.org/) on Oct 26, 2018. The clades for ERV families were obtained from the Dfam database (*97*). Genome mappability along the reference genome was calculated by using the GEM-mappability program from the GEM (GEnome Multitool) suite (*98*).

### Gene Ontology enrichment analysis

GO enrichment analyses for gene lists were performed by using Metascape (*99*). GO enrichment analyses for genomic regions (*e.g*., peaks and enhancers) were performed with GREAT (*100*).

### Epigenetic annotation of regulatory elements

Different types of regulatory elements are defined based on histone modifications and genomic distribution. Promoters-CREs are defined as peaks that are within 500 bp from TSSs, distal-CREs as peaks that are more than 500 bp away from TSSs, and enhancers as H3K27ac peaks that are more than 500 bp away from TSSs.

### Annotation of the genes adjacent to given genomic regions

The genes adjacent to given genomic regions (*e.g*., peaks, enhancers, ERV elements) were determined by comparing their TSSs against specified genomic regions using the *window* function of BEDTools v2.29.2 (*101*). The TSS annotation file was retrieved from the Ensembl database (release 102) using BioMart (*96*).

### TE enrichment analysis

TE enrichment analysis using the TEENA web server we recently developed (*102*). To control for Family-Wise Error Rate, the calculated P-values were further adjusted with the Bonferroni method.

### Interspecies differential expression analysis

Taking the calculated TPM values as input, differential expression analysis between human and mouse was performed by using the ExprX package we previous developed (*56*). DEGs were determined by using the threshold: adjusted p-value < 0.05, |log2foldChange| > 1.

### Statistical analysis and data visualization

All statistical analyses were performed with R statistical programming language (*103*). Heatmaps for ChIP-seq data were generated using DeepTools v3.5.1 (*95*). Heatmaps for gene expression clustering analysis were generated using pheatmap (*104*). Representative tracks for RNA-seq, ChIP-seq, and ATAC-seq data were visualized using IGV v2.13.1 (*105*).

## Acknowledgments

This study utilized the computational resources of Yangzhou University College of Veterinary Medicine High-Performance Computing cluster.

## Funding

National Natural Science Foundation of China 31900422 (MS) Ministry of Science and Technology of China G2023014079L (MS) The 111 Project D18007 (MS)

Priority Academic Program Development of Jiangsu Higher Education Institutions (MS)

## Author contributions

Conceptualization: W.B, M.S

Methodology: C.D, H.F, J.J, S.C, C.L, Y.L, M.L, M.S

Investigation: C.D, H.F, J.J, J.Y, N.V, C.Z, C.L, Y.L, M.S

Visualization: C.D, H.F, J.J, J.Y, N.V, M.S

Supervision: W.B, M.S

Writing—original draft: C.D, H.F, M.S

Writing—review & editing: C.D, H.F, M.L, N.V, W.B, M.S

## Data availability

All the raw and processed sequencing data generated in this study have been submitted to the NCBI Gene Expression Omnibus (GEO; https://www.ncbi.nlm.nih.gov/geo/) under accession GSE266044 and GSE266045. All data needed to evaluate the conclusions in the paper are present in the paper and/or the Supplementary Materials.

## Table of Contents for Supplementary Materials

### Supplementary Materials and Methods

Figs. S1-S15

## Datasets

Data S1: Sources of multi-omics data used in this study

Data S2: Cell-specific CREs annotated for various types of human immune cells

Data S3: GO enrichment for the distal csCREs of various types of human immune cells

Data S4: Enrichment of TF motifs in the CREs for various types of human immune cells

Data S5: TE families enriched in the distal csCREs for various immune cell types

Data S6: Taxonomy of the TE families enriched in at least one type of human immune cells

Data S7: TE families enriched in monocyte enhancers

Data S8: Genes adjacent to SPI1-bound TE-enhancer in human monocytes

Data S9: Motifs enriched in TE-derived monocyte enhancers

Data S10: TE families enriched in the peaks of different TFs in human monocytes

Data S11: Differentially expressed genes after SPI1 knockdown in THP-1 cells

Data S12: Genomic loci with altered H3K27ac levels after SPI1 knockdown

Data S13: Genes adjacent to SPI1-bound loci with decreased H3K27ac after SPI1 knockdown

Data S14: TE families enriched in the lineage-specific enhancers for human and mouse monocytes

Data S15: Differentially expressed genes between human and mouse monocytes

Data S16: TE-derived enhancers adjacent to genes with higher expression in human monocytes

Data S17: The oligos designed for the experiments in this study

