## Supplemental materials for "Unraveling transposable element-mediated regulatory landscape in diverse human immune cells"

**Supplementary Materials and Methods**

**ChIP-seq**

Approximately 1 × $10^7$ cells were used per immunoprecipitation. At first, the cells were fixed with 1% formaldehyde (Sigma-Aldrich, 252549) at room temperature for 10 min. A final concentration of 0.2 M glycine was added to quench the fixation. Nuclear lysates were fragmented by Covaris S220 (Adaptive Focused Acoustics®). For each tissue, 3 µg of H3K27ac (Abcam, ab177178) antibodies were mixed with 25 µL Protein A/G Dynabeads (Selleck. cn, B23201), with rotation at 4°C for 4 h. For per immunoprecipitation, about 50 µg of chromatin was incubated with antibody-bead complexes overnight at 4 °C. ChIP-DNA was purified by MinElute PCR Purification kit (QIAGEN, Cat# 28004). ChIP-seq libraries were constructed using the MGIEasy DNA library prep kit (1000006985, BGI-Shenzhen, China), and were sequenced as 100 bp single-ended reads on BGISEQ-500 platform (BGI-Shenzhen, China).

**RNA-seq**

Trizol reagent (Qiagen) was used for total RNA isolation. The mRNA was purified from total RNA using poly-T oligo-attached magnetic beads, and then the fragmented mRNA was used for the synthesis of cDNA via reverse transcription followed by purification. The PCR-amplified cDNA purification was conducted via an AMPure XP system (Beckman Coulter, Beverly, USA), and the obtained cDNA fragments were 250–300 bp long. The libraries were sequenced as 150 bp paired-end reads on BGISEQ-500 platform (BGI-Shenzhen, China).

**Western blot**

THP-1 cells were lysed with a RIPA buffer supplemented with phosphatase inhibitors for 20 mins on ice, and centrifuged at 12,000 rpm at 4 °C for 15 min. After quantified using a BCA protein assay kit (Solarbio, Beijing, China), equal amounts (20 µg) of denatured proteins were loaded on 10% SDS-PAGE gel for electrophoresis, transferred to PVDF membranes (Immobilon, Darmstadt, Germany), and then blocked with 5% skim milk at room temperature for 1 h. PVDF membranes were incubated overnight at 4°C with anti-SPI1 (1:1,000, sc-390405, SANTA CRUZ BIOTECHNOLOGY, INC.), and then washed with 1×TBST (Solarbio, Beijing, China) before incubation with secondary antibodies (1:5,000). Protein bands were visualized using Enhanced Chemiluminescence (Abbike, BMU102, Wuhan, China) and then exposed with Tanon 4600 (Tianneng, Shanghai, China). HSP90 was used as the internal control.

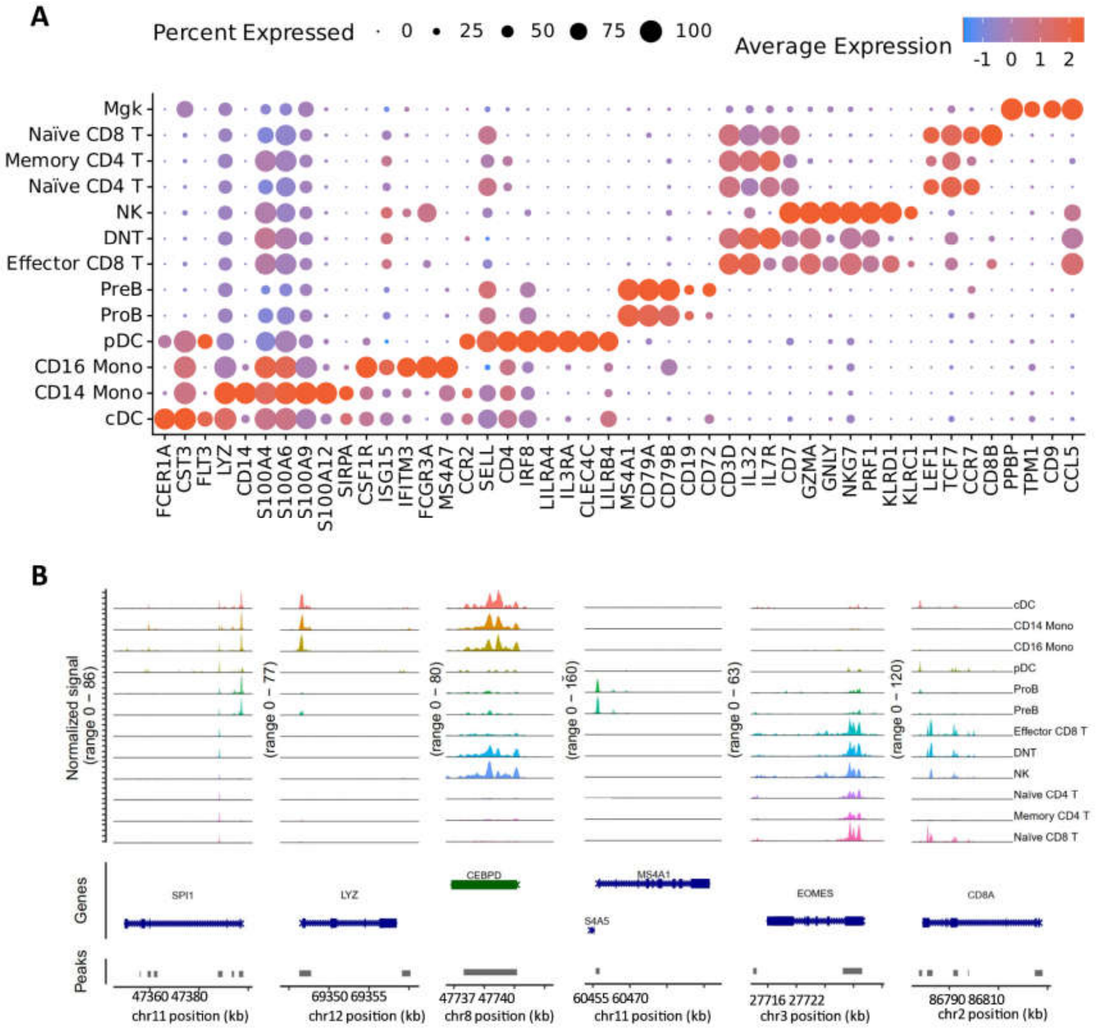


**Fig. S1. Expression and chromatin accessibility of canonical markers across major human immune cell populations**

(**A**) The expression patterns of canonical marker genes across major immune cell populations. (**B**) Chromatin accessibility profiles surrounding canonical marker genes across major immune cell populations. These figures are based on the scRNA-seq and scATAC-seq data of human PBMCs.

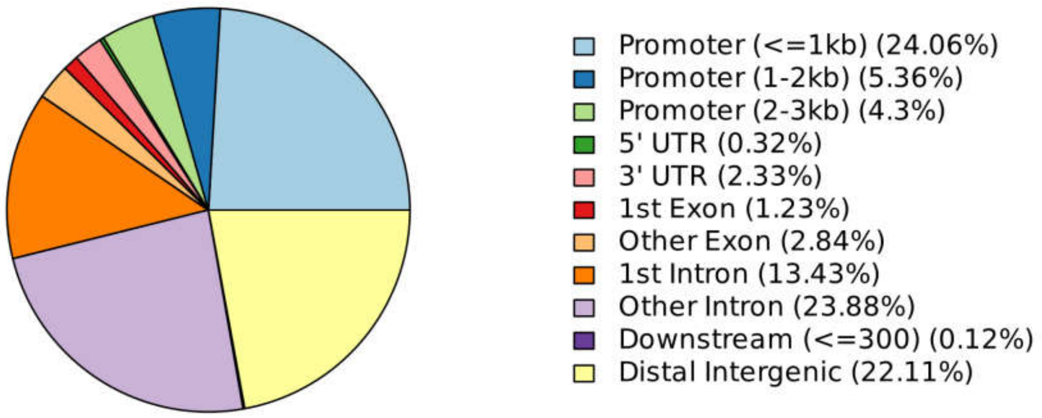


**Fig. S2. Genomic distribution of CREs in major human immune cell populations**

Pie chart of the genomic distribution of CREs, annotated using scATAC-seq data of human PBMCs. The CREs for different immune cell populations are pooled together for analysis. Genomic distribution was annotated by using ChIPseeker.

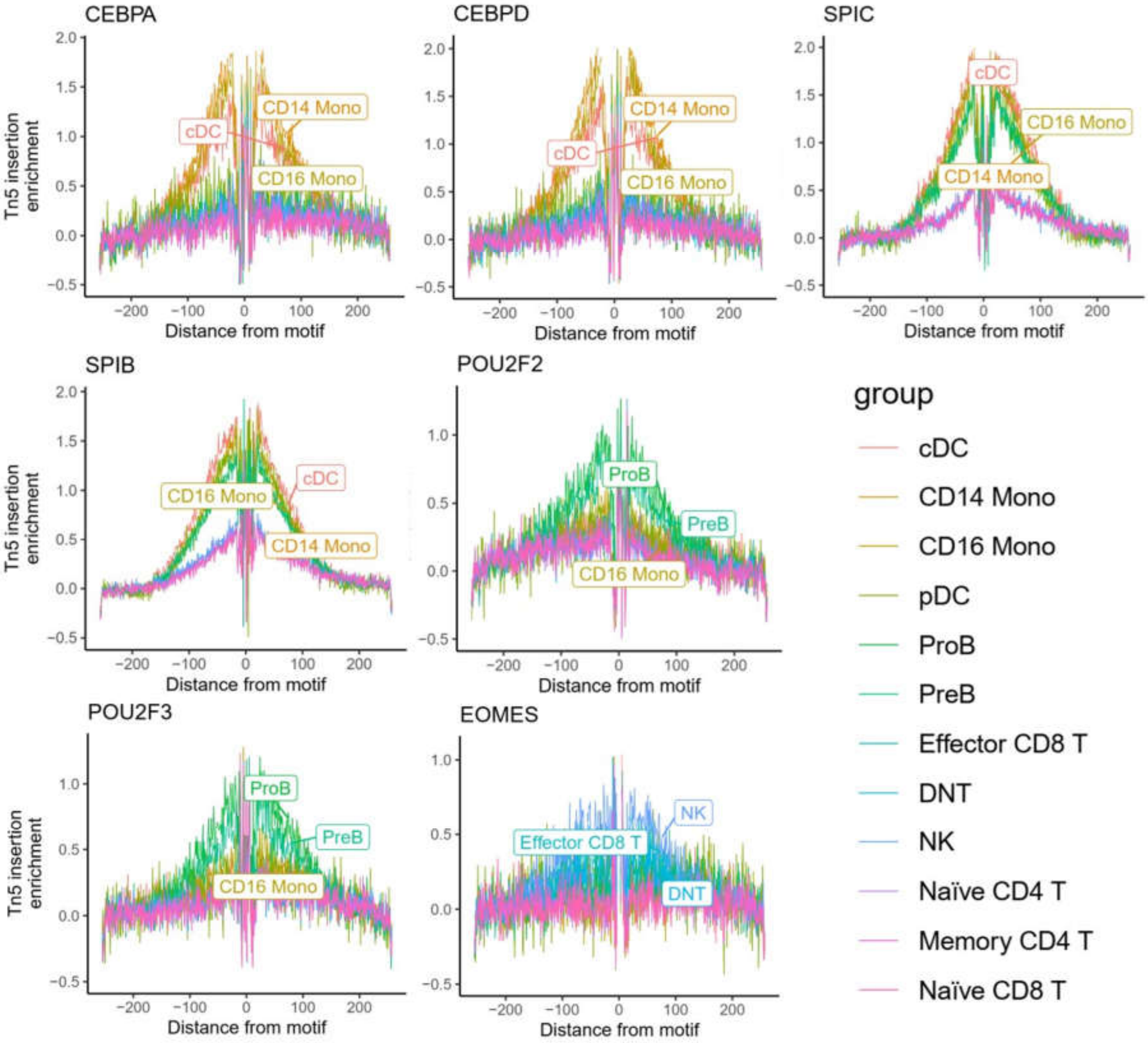


**Fig. S3. TF footprints of selected TF binding sites in major human immune cell populations**

TF footprints for selected binding sites show significant enrichment across various immune cell populations. Footprinting analysis was conducted using JASPAR2020 to incorporate motif information.

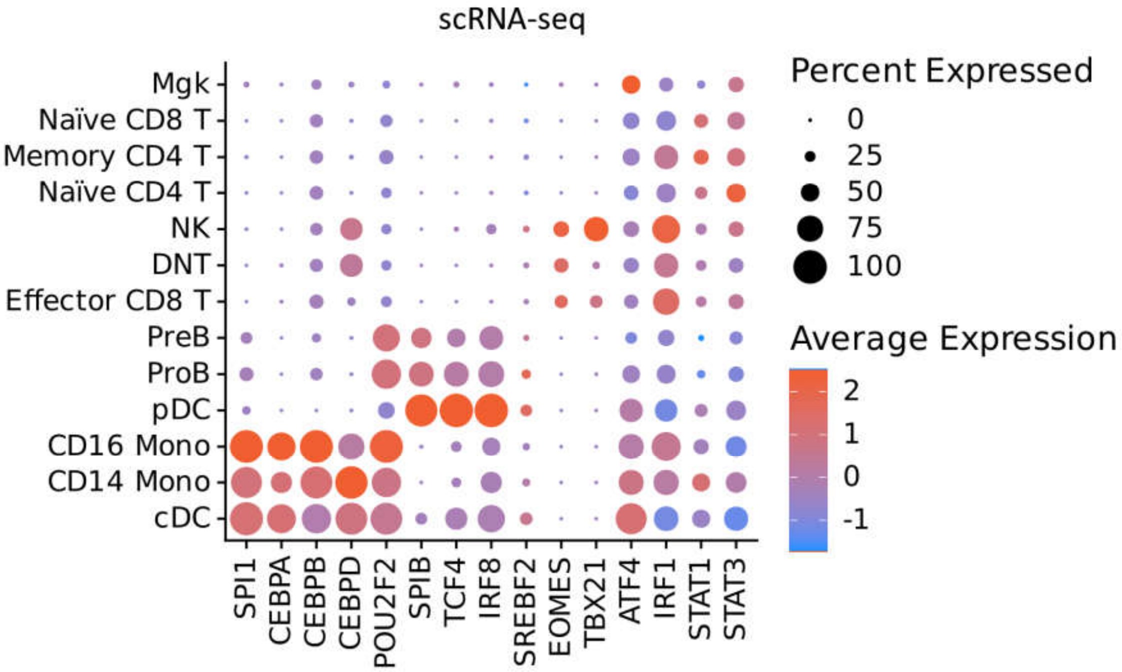


**Fig. S4. Expression of multiple transcription factors in major human immune cell populations**

The dot plot shows the expression patterns of selected cell-specific TFs in major immune cell populations based on scRNA-seq data of human PBMCs.

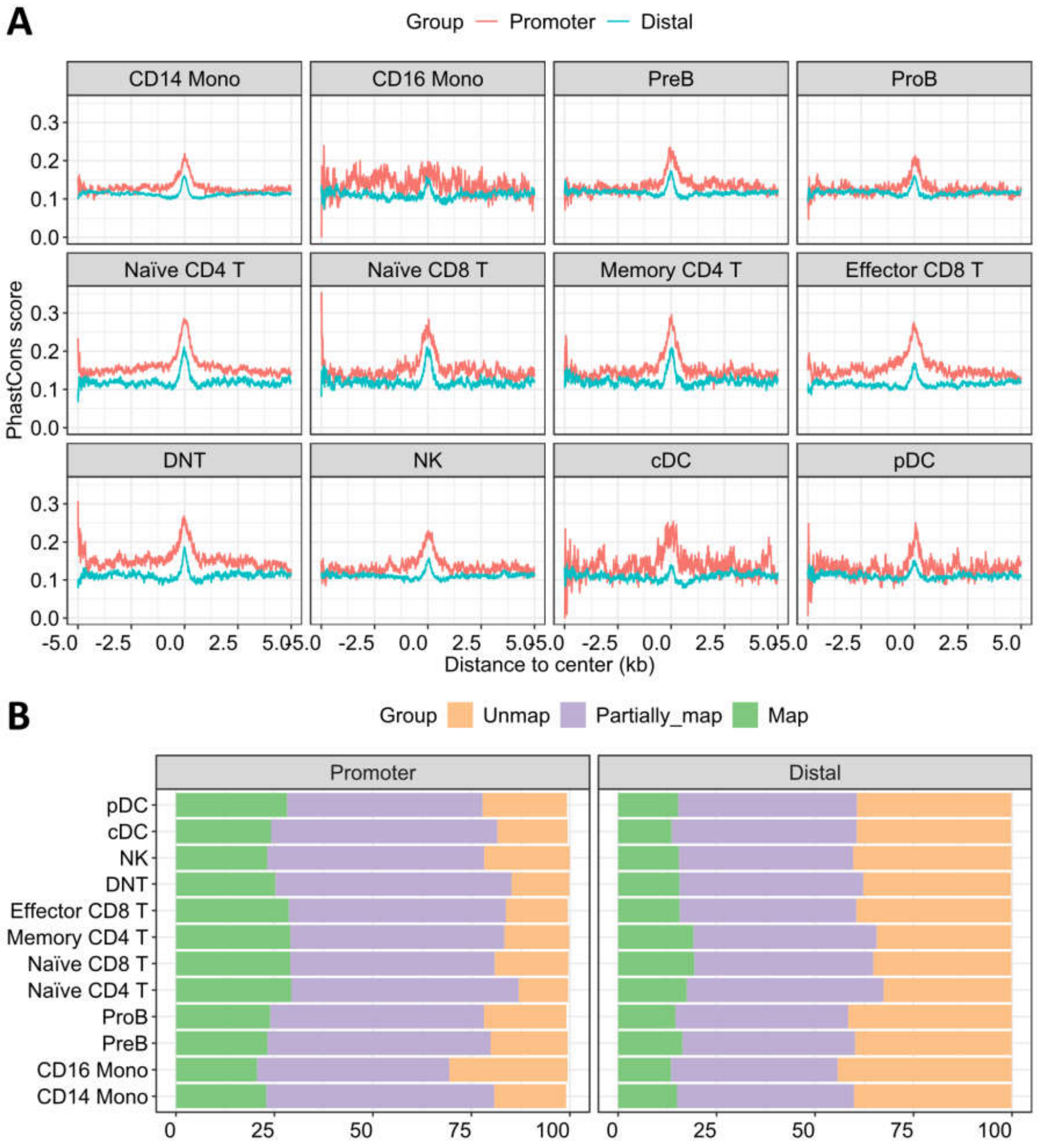


**Fig. S5. Conservation of CREs in major human immune cell populations**

(**A**) The conservation levels of promoter-proximal and distal CREs, as measured by using the UCSC phastCons score. (**B**) The proportions of promoter-proximal and distal CREs for major human immune cell populations that can be mapped to mouse genome or not. The mapping of human CREs to mouse genome was performed by using UCSC liftOver.

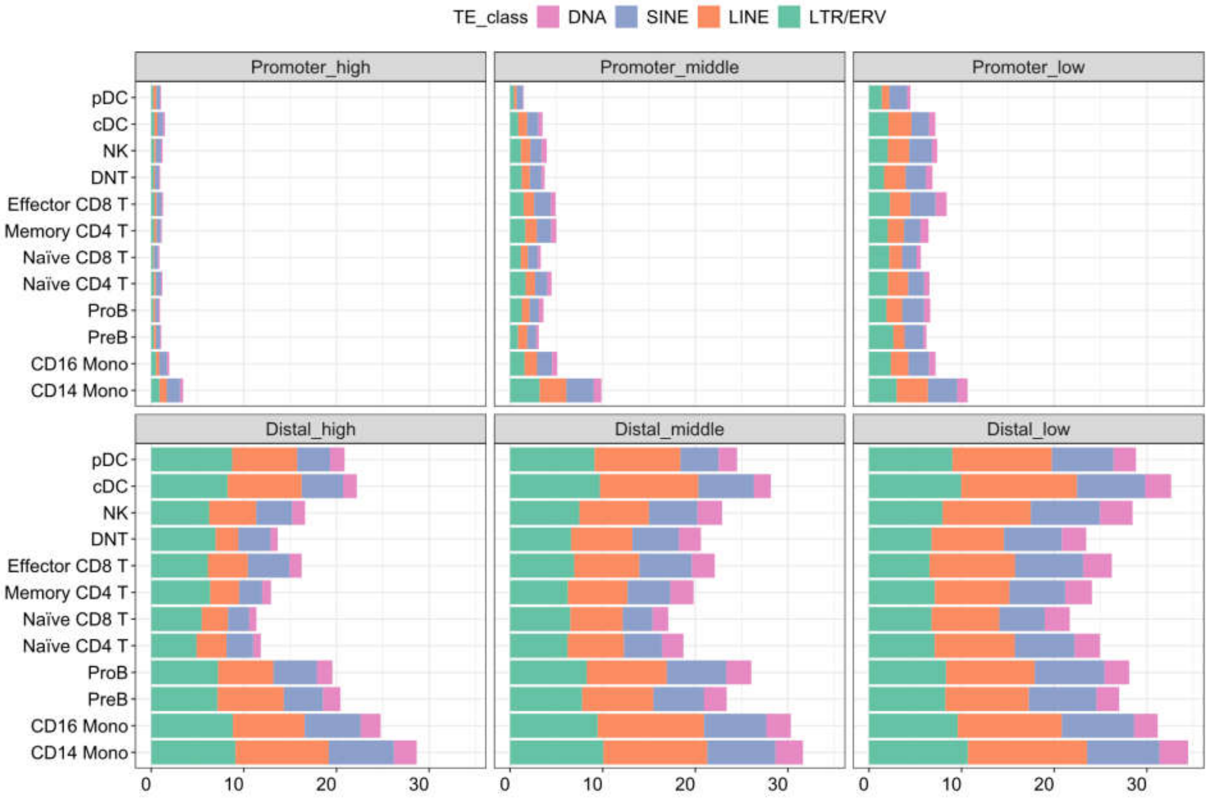


**Fig. S6. Overlap between CREs and TEs in major human immune cell populations**
This figure shows the overlap ratio between immune csCREs and TEs. Promoter-proximal CREs are defined as those within ±500 bp from the TSSs. CREs are classified into three equally-sized groups (high, middle, and low) according to their peak score.

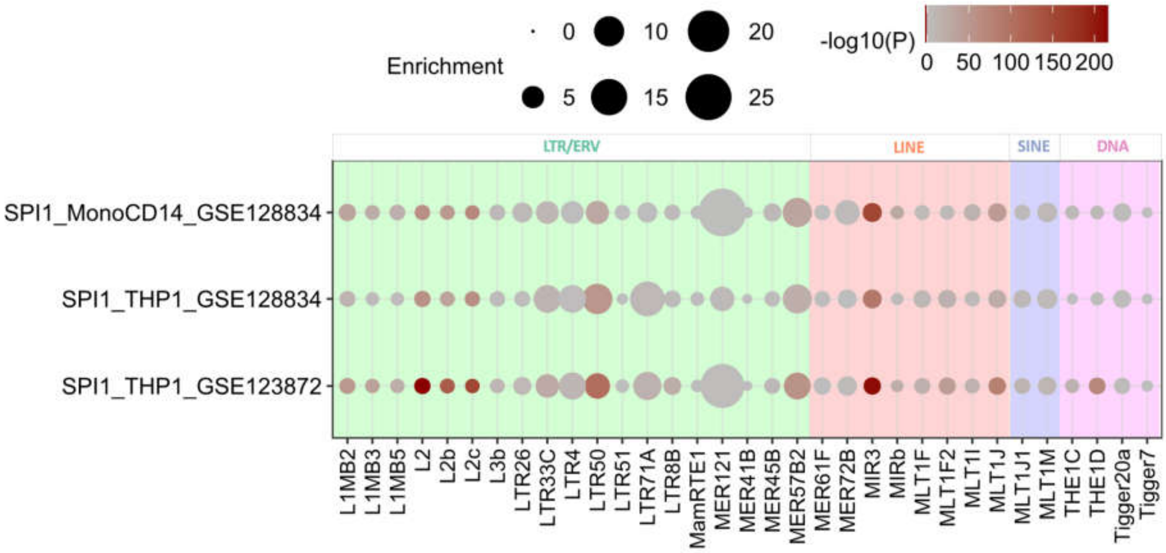


**Fig. S7. TE enrichment patterns of SPI1 peaks in human THP-1 monocyte cell line and primary monocytes**

This figure compares the TE enrichment pattern at the SPI1 binding loci between primary $CD14^+$ monocytes and THP1 monocyte cell line. The TE families significantly enriched in $CD14^+$ monocyte enhancers are included for visualization.

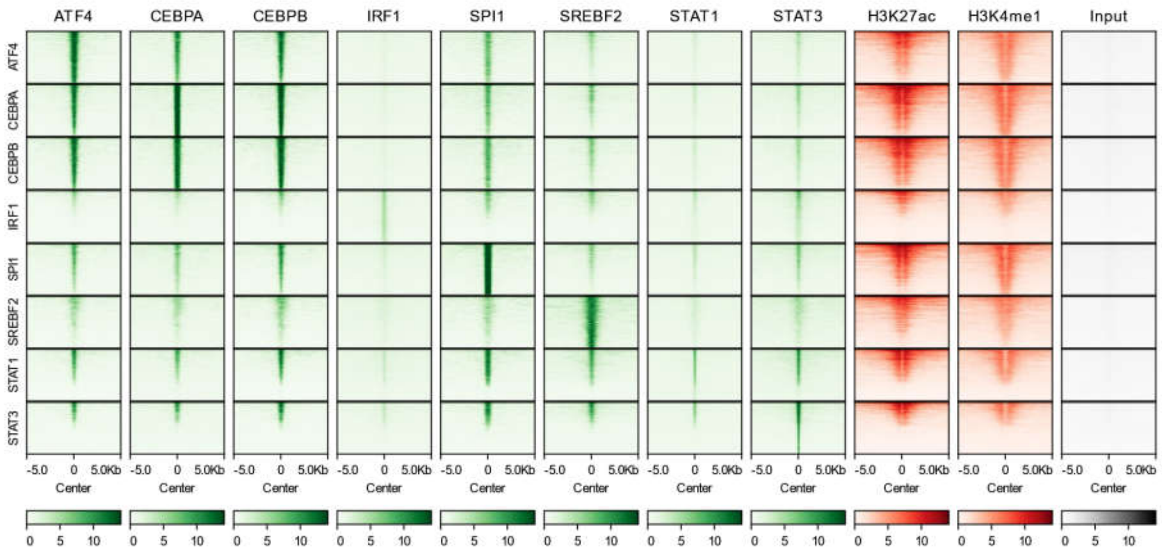


**Fig. S8. Epigenetic characteristics of selected TFs in human monocytes**

The heatmap illustrates the epigenetic characteristics of the binding sites for multiple TFs in human monocytes. This figure is derived from the reanalysis of human monocyte ChIP-seq data (GEO accession: GSE137031, GSE123872, GSE100381, GSE129202, GSE43036, GSE120945) and highlights the top 2000 binding peaks for each TF.

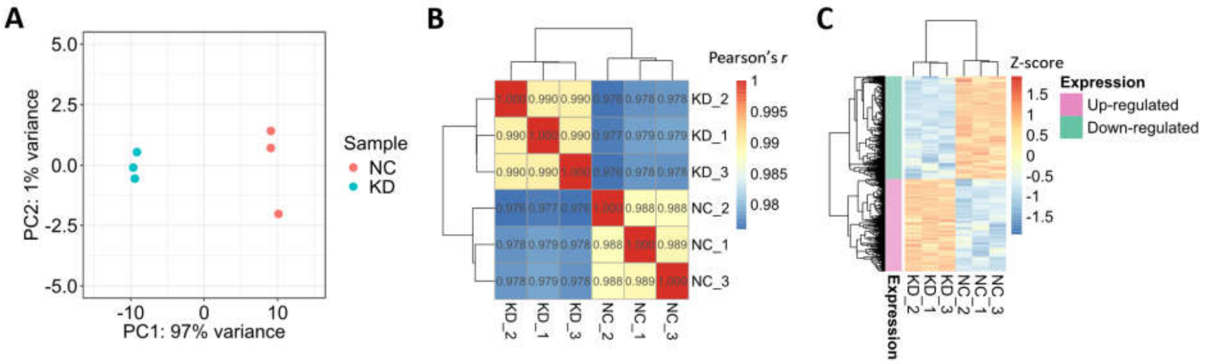


**Fig. S9. Differential expression comparison before and after SPI1 knockdown in THP-1 cells**
(**A**) PCA plot of the relationship among RNA-seq replicates, based on the top 1000 genes with the highest expression variance. (**B**) Correlation heatmap of gene expression relationships based on RNA-seq data. The color gradient represents Pearson's *r*. (**C**) Expression profiles of DEGs identified after SPI1KD. The color gradient represents the Z-score.

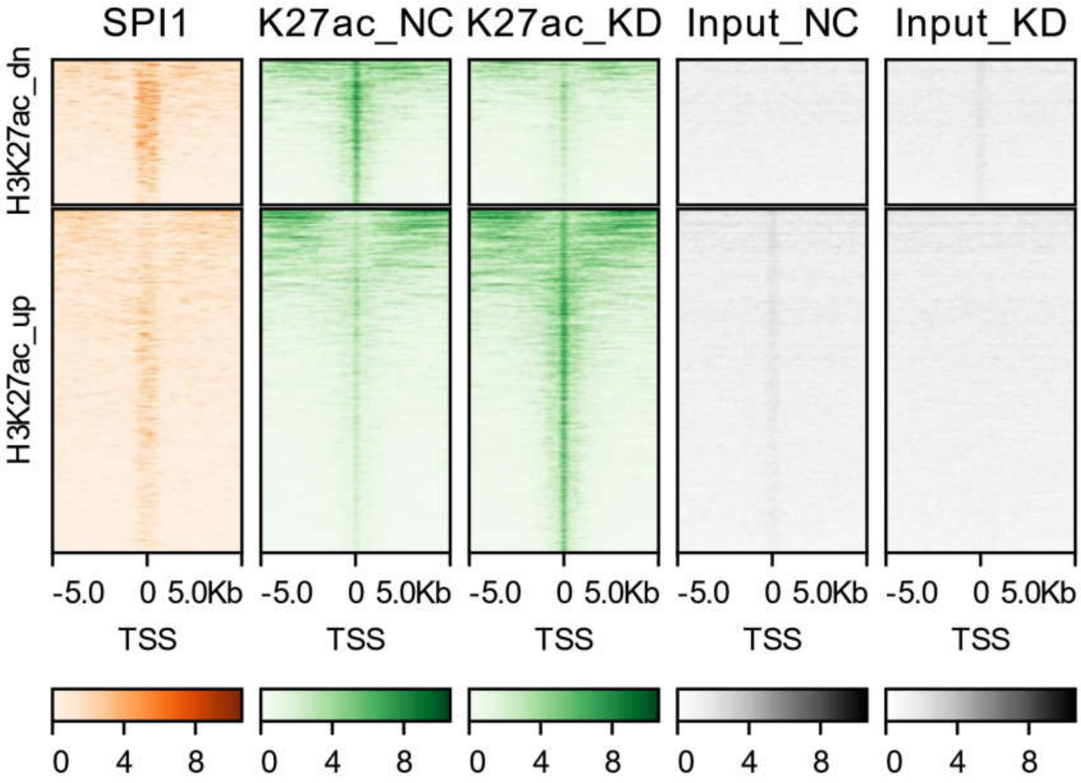


**Fig. S10. SPI1 binding intensity on genomic loci with altered H3K27ac levels after SPI1KD**
The heatmap illustrates a higher SPI1 binding intensity at the genomic loci where H3K27ac level get decreased following SPI1KD.

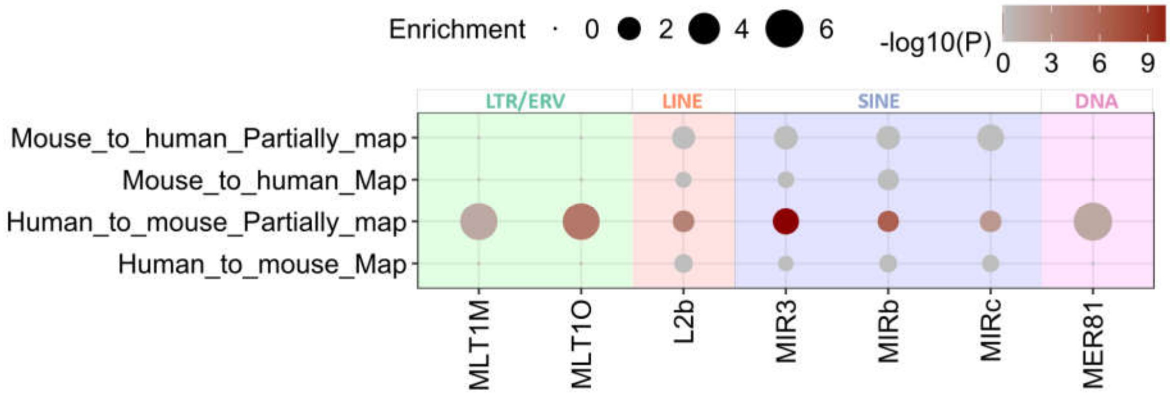


**Fig. S11. TE enrichment patterns in the enhancers shared between human and mouse monocytes**

This figure shows the significant enrichment of TE families in the enhancers shared between human and mouse monocytes (GEO accession: GSE58310, GSE95049). The enhancers entirely or partially mapped between human and mouse are annotated by using UCSC liftOver.

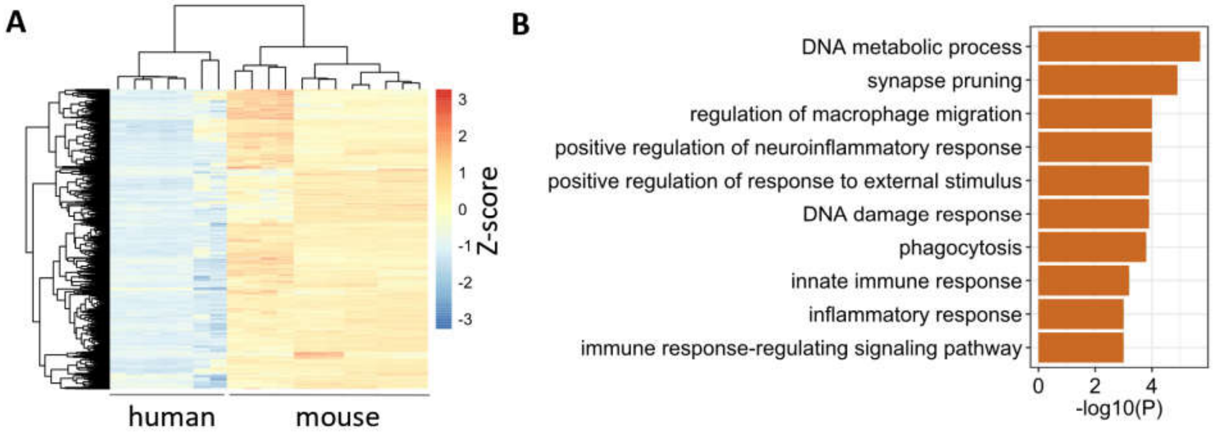


**Fig. S12. Analysis of highly expressed genes in mouse monocytes based on RNA-seq data**
(**A**) Expression profile for the genes with higher expression in mouse monocytes relative to human. This figure is based on RNA-seq data collected from the GEO database, with accessions: GSE118165, GSE115736, GSE95411, GSE217597, GSE117149, GSE116177). (**B**) GO enrichment results of the genes with higher expression in mouse monocytes relative to human.

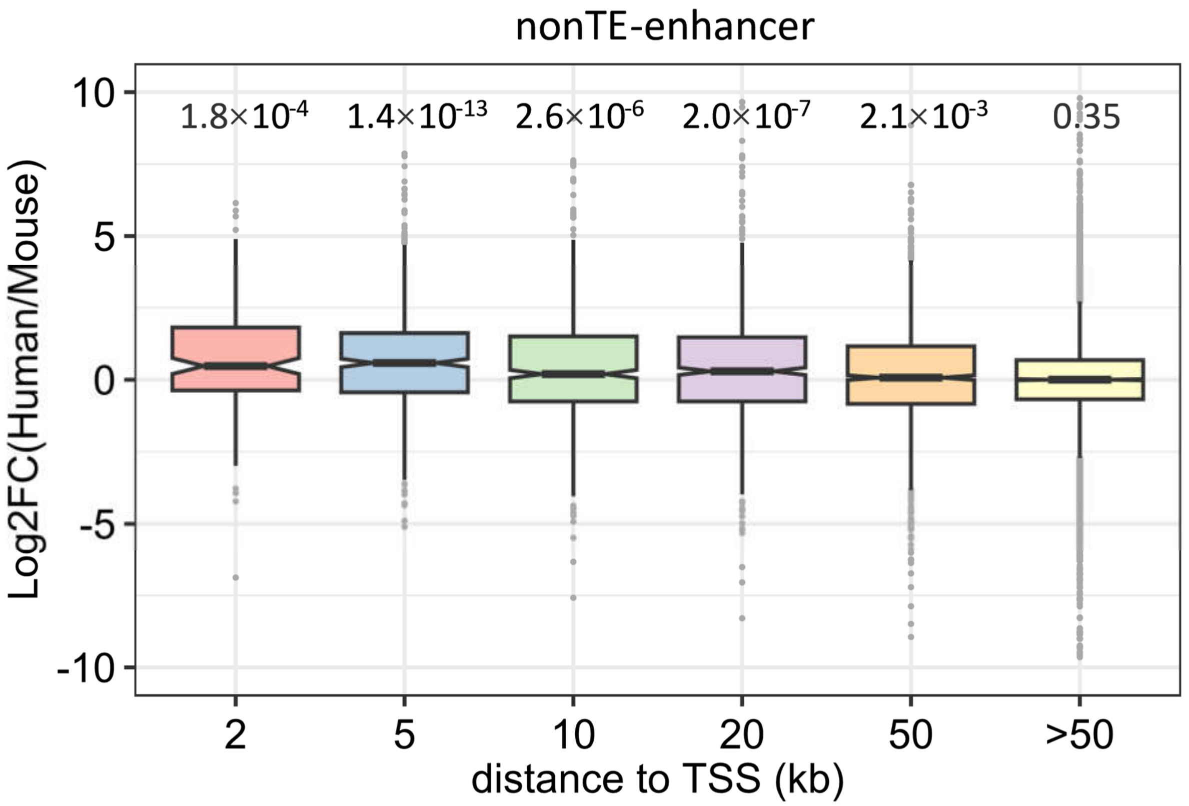


**Fig. S13. Interspecies expression alterations of the genes associated with lineage-specific monocyte enhancers not derived from TEs**

The boxplots compare the log2foldchange of gene expression (human *vs.* mouse) for the genes with human-specific monocyte enhancers occurring at different distance thresholds. The P-values calculated by using Student's t-test again zero are denoted. This figure is related to **Fig. 6F**. It shows that both TE-derived and nonTE-derived lineage-specific enhancers are associated with the altered expression of monocyte genes across species.

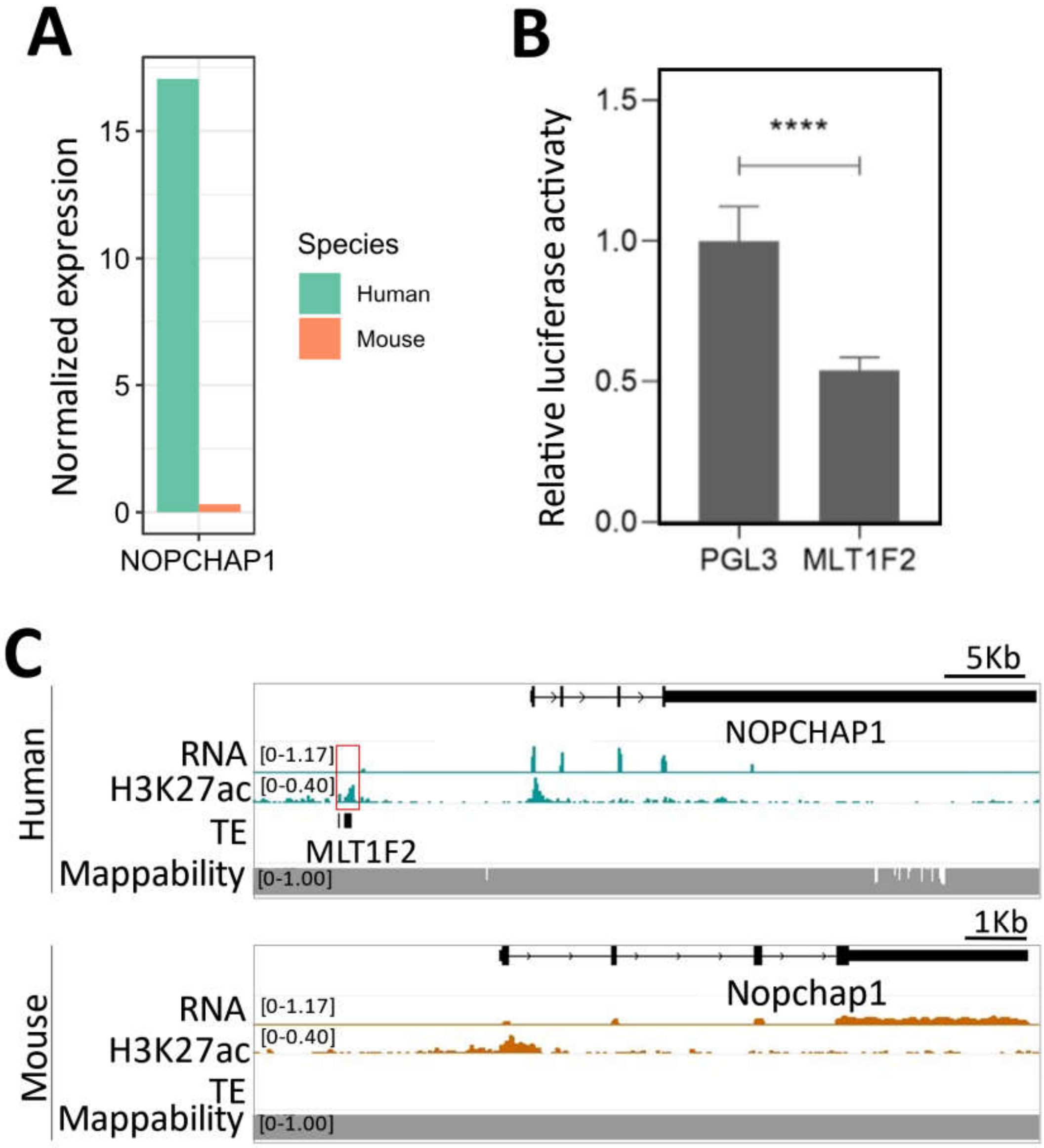


**Fig. S14 Comparative Analysis of NOPCHAP1 Expression and Associated Enhancers in Human and Mouse Monocytes**

(**A**) Expression level of *NOPCHAP1* between human and mouse monocytes. The expression level was measured as normalized TPM values. (**B**) IGV tracks show the transcriptomic and epigenomic pattern flanking *NOPCHAP1* which is adjacent to putative lineage-specific TE-enhancers. (**C**) Luciferase reporter assay results for the TE elements associated with the putative lineage-specific enhancers adjacent to *NOPCHAP1*.

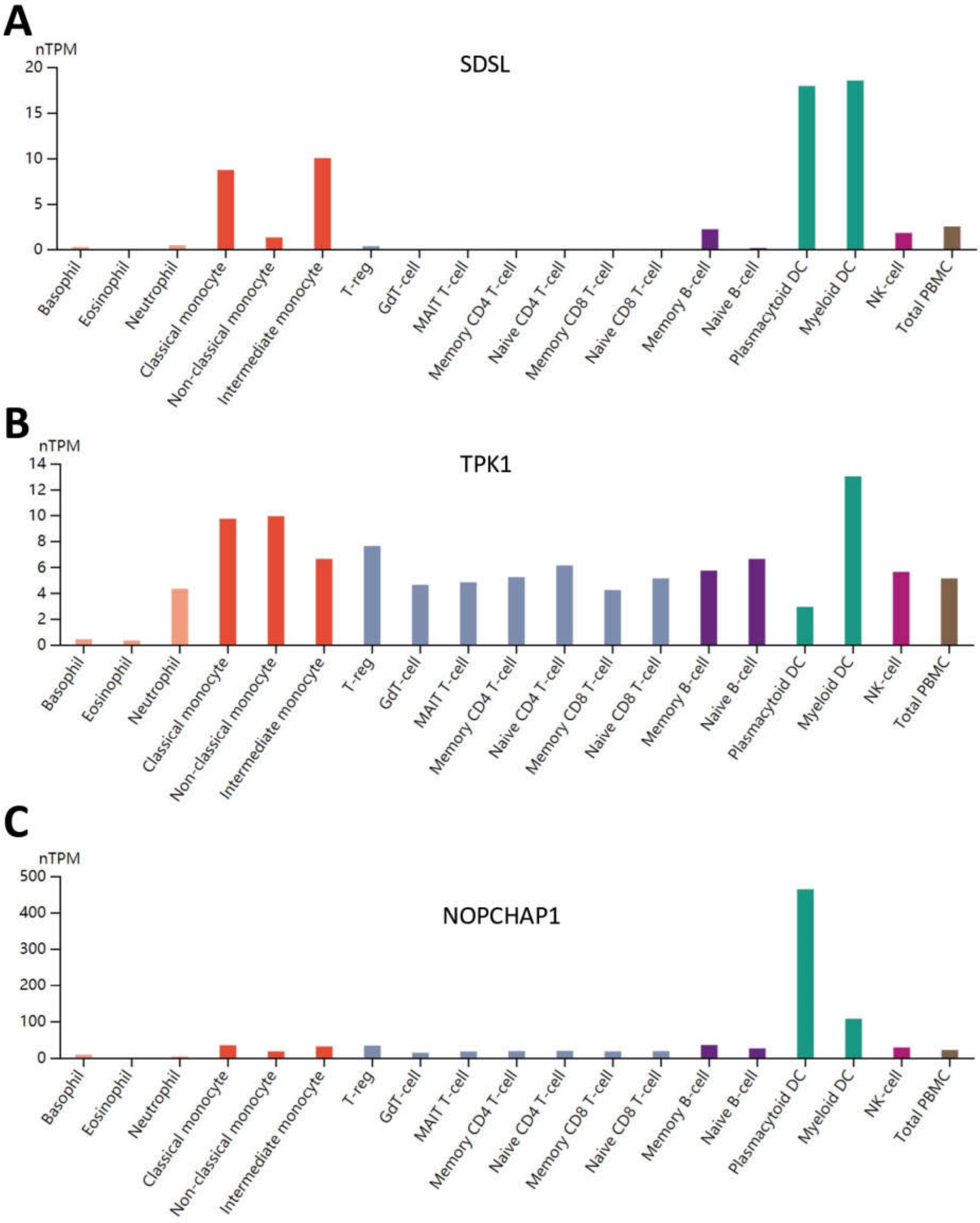


**Fig. S15 Comparative of *SDSL*, *TPK1*, and *NOPCHAP1* Expression in Human Immune Cells.** The expression profiles of *SDSL*, *TPK1,* and *NOPCHAP1* in different types of immune cells. This figure is based on the data from Human Protein Atlas database (https://www.proteinatlas.org/).
